# Shedding Light on the Threespine Stickleback Circadian Clock

**DOI:** 10.1101/2021.06.01.446630

**Authors:** Marie-Pier Brochu, Nadia Aubin-Horth

## Abstract

The circadian clock is an internal timekeeping system shared by most organisms, and knowledge about its functional importance and evolution in natural environments is still needed. Here, we investigated the circadian clock of wild-caught threespine sticklebacks (*Gasterosteus aculeatus*) at the behavioural and molecular levels. While their behaviour, ecology, and evolution are well studied, information on their circadian rhythms are scarce. We quantified the daily locomotor activity rhythm under a light-dark cycle (LD) and under constant darkness (DD). Under LD, all fish exhibited significant daily rhythmicity, while under DD, only 18% of individuals remained rhythmic. This interindividual variation suggests that the circadian clock controls activity only in certain individuals. Moreover, under LD, some fish were almost exclusively nocturnal, while others were active around the clock. Furthermore, the most nocturnal fish were also the least active. These results suggest that light masks activity more strongly in some individuals than others. Finally, we quantified the expression of five clock genes in the brain of sticklebacks under DD using qPCR. We did not detect circadian rhythmicity, which could either indicate that the clock molecular oscillator is highly light-dependent, or that there was an oscillation but that we were unable to detect it. Overall, our study suggests that a strong circadian control on behavioural rhythms may not necessarily be advantageous in a natural population of sticklebacks and that the daily phase of activity varies greatly between individuals because of a differential masking effect of light.

**Summary statement:** We found that in wild-caught threespine sticklebacks, the circadian clock does not control locomotor activity in most, but not all, individuals. Sticklebacks are mostly nocturnal, although interindividual variation exists.

## INTRODUCTION

Many behaviours and physiological processes in living organisms exhibit daily rhythmicity, for example: locomotor and feeding activity, hormone secretion, and metabolism (Refinetti, 2016). Some of these rhythms persist in the absence of external cues, because they are driven by an endogenous mechanism called the circadian clock (Kumar, 2017). Found in almost all life forms, this internal clock usually has an intrinsic period of approximately 24 h and is entrained by temporal signals such as the light-dark cycle, so that the phase of circadian rhythms is synchronized with relevant environmental variables (ex.: being awake when feeding or mating opportunities are present). The circadian clock thus allows the anticipation of daily environmental changes and the coordination of biological functions, and can have fitness consequences (Vaze and Sharma, 2013; Dominoni et al., 2017). The heart of the circadian clock is a cell-autonomous molecular oscillator made up of a transcription-translation feedback loop that involves positive and negative elements (Bell-Pedersen et al., 2005). In mammals, BMAL and CLOCK are positive elements that induce the transcription of *period* (*per*) and *cryptochrome* (*cry*). PER and CRY are negative elements that inhibit their own transcription by down-regulating the activity of BMAL and CLOCK (Rosensweig and Green, 2020). Generally speaking, the expression level of *bmal* and *clock* is in antiphase with that of *per* and *cry* (Takahashi, 2017). These four clock genes are highly conserved in animals, but, because of genome duplication events, several of them retain two paralogous copies in the different vertebrate lineages (Bell-Pedersen et al., 2005).

In the last decade, our knowledge of the organization and functioning of circadian rhythms in animals has expanded with the study of various wild species, building on the work mostly acquired in laboratory settings with model organisms (*Mus musculus, Danio rerio, Drosophila melanogaster*) (Kronfeld-Schor et al., 2013; Schwartz et al., 2017). This growing body of research shows that the implication of the circadian clock in driving biological rhythms can vary greatly depending on a species’ biology (reviewed in Bloch et al., 2013; Hazlerigg and Tyler, 2019) and that the opportunities, threats and challenges that organisms face in natural environments can influence their circadian rhythms (Hut et al., 2012; Helm et al., 2017). For example, some species adjust the phase of their circadian activity rhythm in response to light intensity (Chiesa et al., 2010), food availability (Lopez-Olmeda et al., 2010; Ware et al., 2012), predation risk (Pellman et al., 2015) and social interactions (Fuchikawa et al., 2016). In order to improve our understanding of the functional importance of the circadian clock in nature (i.e. the benefits it provides to an individual in a given environment) and which selection pressures can shape the evolution of circadian rhythms, we must continue to investigate a diversity of species that have evolved in various ecological contexts and that are amenable to experimental and physiological studies (Kronfeld-Schor et al., 2013; Schwartz et al., 2017). The threespine stickleback (*Gasterosteus aculeatus*) is well studied in ecology and evolution (McKinnon et al., 2019). This small fish is also well suited to answer questions about the ecological and evolutionary implications of the circadian clock through the study of its natural phenotypic variation, which can be combined with experimental work. Stickleback ecotypes are found in diverse habitats (marine waters, salt marshes, streams, rivers, lakes, etc.) and display morphological, physiological, and behavioural adaptations to these environments (Bell and Foster, 1994; Ostlund-Nilsson et al., 2007; Kitano et al., 2010; Di-Poi et al., 2014; Di Poi et al., 2016; Ishikawa et al., 2019). Many of the environmental pressures that differ between ecotypes such as the presence of predators and parasites, prey availability, light intensity and social interactions (Ostlund-Nilsson et al., 2007) have the potential to influence circadian rhythms (Helm et al., 2017). This could be achieved either through selective pressure resulting in genetic divergence, or through phenotypic plasticity, i.e. the effects of the environment on the development of a phenotype, here the circadian rhythm itself. As sticklebacks are also known for their interindividual variation in behaviour, called personality (activity, boldness, sociality, etc. (Huntingford, 1976; Bell, 2005; Wark et al., 2011)), it is also possible that they exhibit interindividual variation in circadian rhythms. So far, it has been suggested that circadian molecular mechanisms may vary between ecotypes similarly to traits at other levels of biological organization, although the functional impact of this difference is not known. For example, using common garden-raised sticklebacks from two lake-stream pairs, a previous study reported that a gene that is part of the molecular oscillator (*cry1ab*) was upregulated in the liver of stream sticklebacks compared to lake ones (Hanson et al., 2017). Studying circadian rhythms in sticklebacks will help us to better understand the functional importance and the evolution of the circadian clock in natural environments.

In comparison to what is known about the ecology and evolution of sticklebacks, very little knowledge is available on their circadian rhythms and clock. In fact, the existence of a circadian clock has never been demonstrated in this species. At the behavioural level, sticklebacks have, to our knowledge, only been studied once under constant light conditions. This study showed that the frequency with which males visited their nests (in the hope of finding eggs deposited by a female) did not display circadian rhythmicity in constant light (Sevenster et al., 1995). Regarding the daily activity rhythm (i.e. under a light-dark cycle), some evidence suggests that sticklebacks are diurnal. For instance, stickleback visual opsins (Rennison et al., 2012) correspond to those of diurnal fish (Carleton et al., 2020). Moreover, previous studies reported that sticklebacks were mostly captured during the day in the wild (Worgan and FitzGerald, 1981; Sjoberg, 1985; Reebs et al., 1995). On the other hand, night activity (Reebs et al., 1984; Quinn et al., 2012) and night feeding (Mussen and Peeke, 2001) have been observed in some sticklebacks. At the physiological level, we know that melatonin levels (a hormone that plays a key role in the regulation of circadian rhythms) are higher during the night than during the day in sticklebacks (Mayer et al., 1997; Kulczykowska et al., 2017; Pomianowski et al., 2020) as in most vertebrates (Challet, 2007; Falcón et al., 2009), but we do not know if this rhythm is driven by the clock or solely by light (Falcón et al., 2009). At the molecular level, time-of-day variation in the expression of *per1b* and *clock1b* has been observed in the liver of sticklebacks, but since this was measured under a light-dark cycle, we do not know if this rhythm is self-sustained (Prokkola et al., 2015).

In this study, using wild-caught threespine sticklebacks, we investigated the circadian clock of this species at the behavioural and molecular levels. Our first objective was to determine if the daily rhythm of locomotor activity is under circadian clock control, and we hypothesized that it is indeed the case. Our prediction was that sticklebacks would show a significant rhythm of locomotor activity under constant darkness (DD). Our second objective was to determine the phase of activity of sticklebacks under LD. We hypothesized that sticklebacks are diurnal. Our prediction was that the daily activity would be mainly performed during the light phase. Our third objective was to quantify the molecular oscillation of five clock genes (*bmal1a, clock1b, clock2, per1b* and *cry1b*) in the brain, an organ that is potentially implicated in the control of circadian rhythms. We hypothesized that clock gene expression shows circadian rhythmicity under DD. Our prediction was that the expression level of *bmal1a, clock1b* and *clock2* would be in antiphase with that of *per1b* and *cry1b*.

## MATERIALS AND METHODS

### Fish sampling and housing

We collected threespine sticklebacks (*Gasterosteus aculeatus*) from the wild population of the lac Témiscouata (47°48’37.1”N 68°51’56.6”W, Québec, Canada) in June 2019. We did not have specific information on the daily activity patterns of this species in the lac Témiscouata. We thus sampled fish with a beach seine so that we could collect all individuals in the water column no matter if they were resting at the bottom of the lake or swimming at the surface. We sampled fish in the morning (around 8:00), in the afternoon (around 15:00) and in the evening (around 19:00) to account for the possibility that some individuals migrate daily between different parts of the lake. Sticklebacks were brought back to the Laboratoire de Recherches en Sciences Environnementales et Médicales (LARSEM) at Université Laval (Québec, Canada). In the animal facility, fish were held in two 1000 L water tanks (n=140/tank) and were fed brine shrimp and nutritious flakes twice a day (morning and late afternoon). They were exposed to non-breeding environmental conditions, a water temperature of 14°C and a 12 h light:12 h dark cycle with lights on at 6:00 and lights off at 18:00.

### Activity monitoring system

To monitor locomotor activity, 18 fish were transferred in an adjacent room and individually placed in 2 L experimental tanks. A white plexiglass separated each tank to prevent fish from seeing each other. Lightning was provided by three full-spectrum LED light bars (Plant 3.0, Fluval) mounted above the tanks. Illuminance was measured by a lux meter (LX1330B, Dr.meter) and was around 500 lux at the water surface. We chose this illuminance value based on previous studies in other fish species (Iigo and Tabata, 1996; Whitmore et al., 2000; Bayarri et al., 2004; Lopez-Olmeda et al., 2010). A dark plastic curtain was hanging in front of the tanks to ensure a constant illumination (or darkness) when we needed to enter the room for maintenance.

Each experimental tank was equipped with an infrared photoelectric sensor (E3Z-D67, Omron) placed in the lower third of the front wall (Fig. S1). We had previously established that this position was optimal to record stickleback movements (Fig. S2). Every time a fish interrupted the infrared light beam that was emitted by the sensor, an output signal was sent to a controller (ILC 131 ETH, Phoenix Contact). Each interruption was counted as one movement. Data was retrieved by connecting a computer to the controller.

### Experimental design

All experimental procedures were approved by the Comité de Protection des Animaux de l’Université Laval (CPAUL 2018066-2). Since we could monitor 18 fish at a time, we divided individuals into three groups (Fig. 1). Individuals were allowed to acclimate to the experimental tanks for at least three days before the start of the experiment. For all three groups, food was provided by hand once a day at random time (previously determined using the RAND() function in Excel software). We used a dim red light when food was provided during the dark phase.

**Fig. 1.**
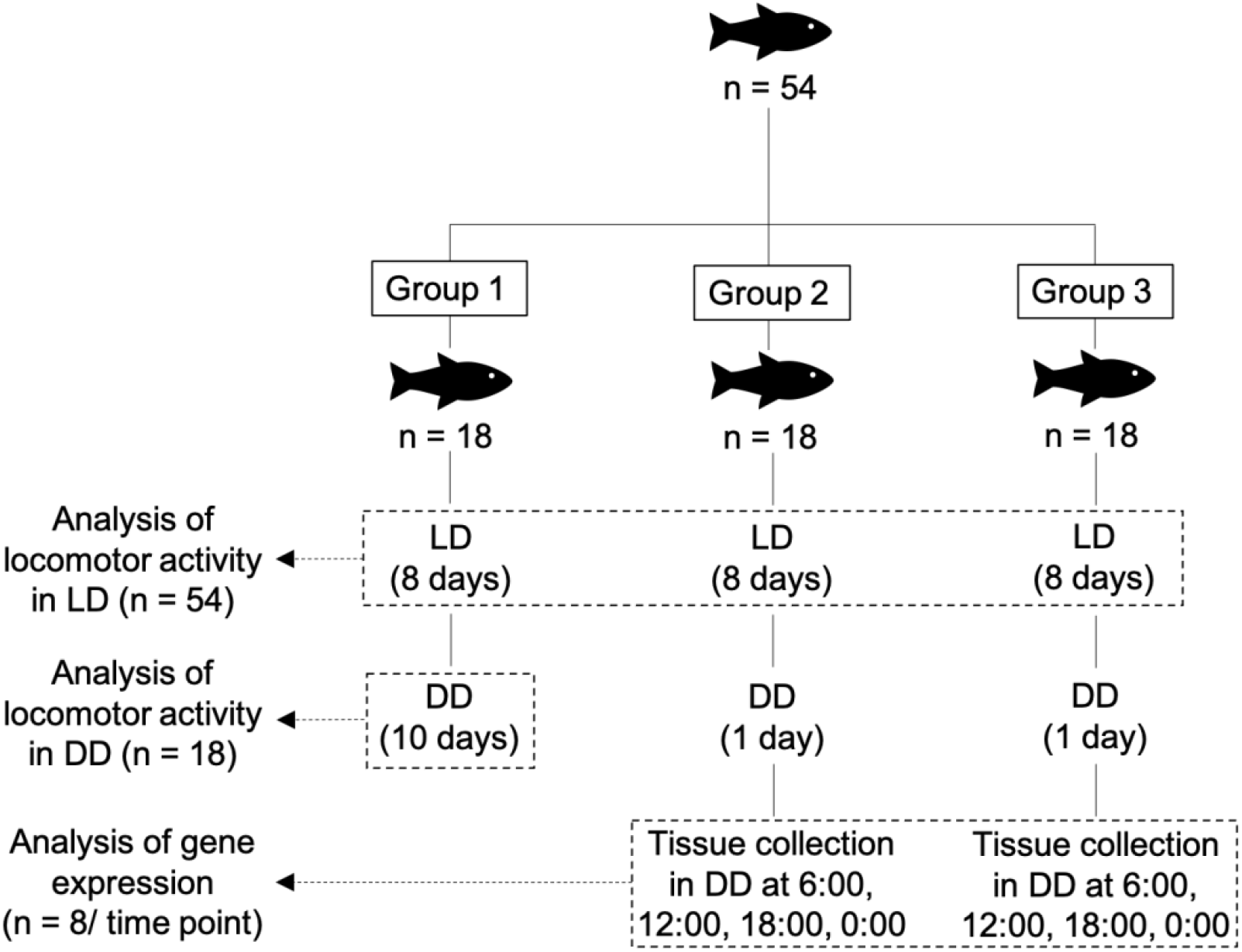
Experimental design. Group 1 was used to quantify locomotor activity under a 12 h light:12 h dark cycle (LD, lights on at 6:00 and lights off at 18:00) and under constant darkness (DD). To that end, group 1 was exposed to LD for 8 days, then to DD for 10 days. Groups 2 and 3 were used to quantify locomotor activity under LD and brain gene expression under DD. Groups 2 and 3 were thus exposed to LD for 8 days, then to DD for 1 day (day 9). The day following the switch to DD (day 10), we sampled the brain of four randomly selected individuals every 6 h throughout a 24-h cycle (6:00, 12:00, 18:00, 0:00).

Group 1 was exposed to a 12 h light:12 h dark (LD) cycle for eight days (lights on at 6:00 and lights off at 18:00) followed by ten days of constant darkness (DD). Group 1 was used to quantify locomotor activity under LD and DD. Groups 2 and 3 were also exposed to LD for eight days and used to quantify locomotor activity under LD. On the ninth day of the experiment with groups 2 and 3, lights were not turned on at 6:00 so all fish were exposed to DD for at least 24 h. On the tenth day, we sampled the brain and the caudal fin of four randomly selected individuals every 6 h throughout a 24-h cycle (6:00, 12:00, 18:00, 0:00), see Fig. 1. Tissue collection was performed in darkness with the help of a dim red light and took less than 3 minutes per fish. After dissection, brains and caudal fins were immediately stabilized in RNA*later* (Ambion) and stored at −20°C. We used caudal fins to determine sex with the IDH genetic sex marker (Peichel et al., 2004).

### Choice of genes

We chose to quantify the expression of *bmal1a, clock1b, clock2, per1b* and *cry1b* although sticklebacks have several other clock genes (Table 1). We chose these five genes for three reasons. First, we wanted to quantify positive (*bmal, clock*) and negative (*per, cry*) elements to have an overall view of the transcription-translation feedback loop. Second, we chose genes that have an ortholog in the zebrafish to compare our results with what is known from this model organism (Table 1). Third, we avoided quantifying *per2a* and *cry1aa* because these two genes are mainly light-induced (in opposition to being clock controlled) in the zebrafish (Pando et al., 2001; Tamai et al., 2007; Vatine et al., 2009), so their expression rhythm rapidly loses its amplitude under DD (ex.: Beale et al., 2013) and thus would not be informative in our study in DD.

**Table 1.** The four core genes of the transcription-translation feedback loop of the clock molecular oscillator in mammals, zebrafish and sticklebacks. The five stickleback genes that we investigated in this study are in bold in the table.

| Gene | Mammals | Zebrafish | Stickleback | Reference for the phylogenetic analysis |
| --- | --- | --- | --- | --- |
| *bmal* | *bmal1* | *bmal1a* | ***bmal1a*** | Wang (2009) |
|  |  | *bmal1b* | - |  |
|  | *bmal2* | *bmal2a* | - |  |
|  |  | - | *bmal2b* |  |
| *clock* | *clock* | *clock1a* | - | Wang (2008b) |
|  |  | *clock1b* | ***clock1b*** |  |
|  | *npas2* | *clock2* | ***clock2*** |  |
| *period (per)* | *per1* | *per1a* | - | Wang (2008a) |
|  |  | *per1b* | ***per1b*** |  |
|  | *per2* | *per2a* | *per2a* |  |
|  |  | - | *per2b* |  |
|  | *per3* | *per3* | - |  |
| *cryptochrome (cry)* | *cry1* | *cry1aa* | *cry1aa* | Liu et al. (2015) |
|  |  | *cry1ab* | *cry1ab* |  |
|  |  | *cry1ba* | ***cry1ba*** |  |
|  |  | *cry1bb* | - |  |
|  | *cry2* | *cry2* | *cry2* |  |
|  | - | *cry3* | - |  |

### Gene expression in the brain

We studied clock gene expression using a quantitative real-time PCR (qPCR) approach. We extracted total RNA in the brain of sticklebacks and performed a DNase digestion using the miRNeasy Mini Kit (Qiagen) combined with the RNase-Free DNase Set (Qiagen). We stored RNA at −70°C. We quantified RNA using the Quant-iT RiboGreen RNA Assay Kit (Invitrogen) and assessed RNA quality and integrity with the RNA 6000 Nano Kit (Agilent). All samples showed RNA integrity numbers (RIN) greater than 9.0. For all samples we reverse-transcribed 10 µL of RNA at 100 ng µL^-1^ with 4 µL of the 5X qScript cDNA SuperMix (Quantabio) and 6 µL of RNase-free water in a final volume of 20 µL. Following the manufacturer’s protocol, thermocycling parameters were 25 °C for 5 min, 42°C for 30 min and 85°C for 5 min.

We obtained cDNA sequences of *bmal1a, clock1b, clock2, per1b* and *cry1b* from the *Ensembl Genome Browser* (version 98) and designed primers using Primer3 (Table 2). We did *in silico* specificity screen with the Amplify4 software to ensure that primers for a given gene were not amplifying any paralogs. We also verified specificity of primers and absence of primer dimers with melting curves (60-95°C). To further guarantee that the primers were amplifying the targeted genes, we analyzed amplicons by Sanger sequencing. We assessed PCR amplification efficiency of each primer pair with a qPCR experiment using a four or five-point standard curve made of a fivefold dilution series of pooled cDNA samples. Efficiency is reported in Table 2.

**Table 2.**
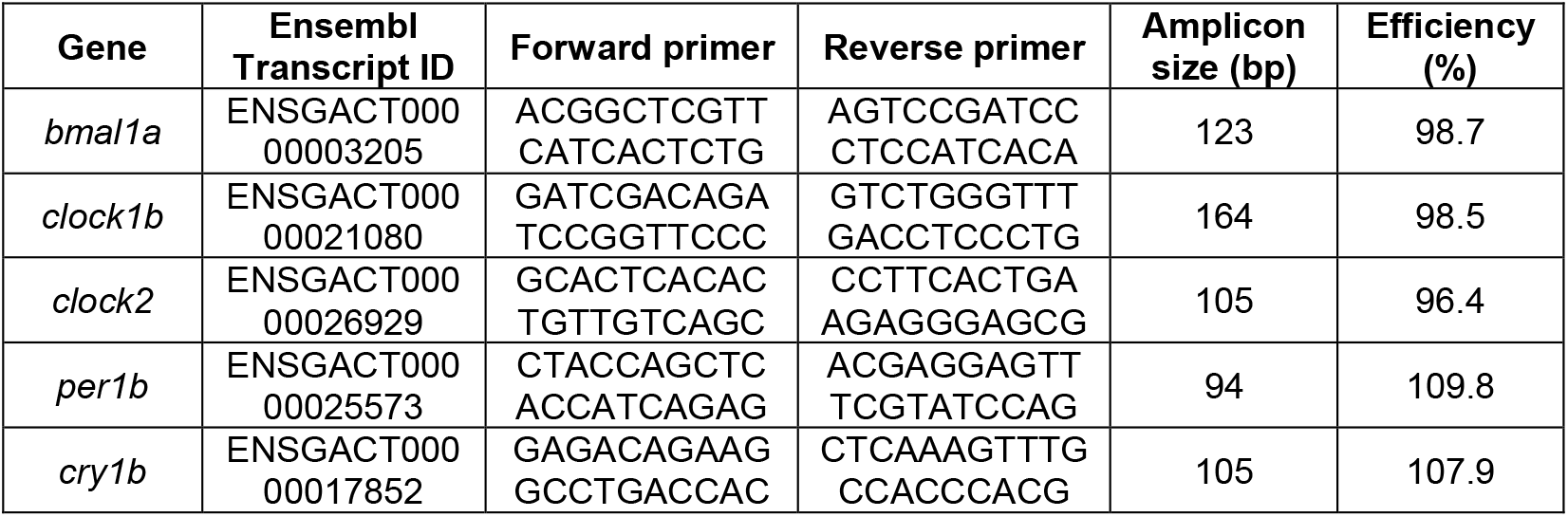
Characteristics of the primers used to quantify clock gene expression. Efficiency (E) was calculated using E = (10 ^-1/slope^ −1)*100 (Pfaffl, 2001).

We performed qPCR experiments in the 7500 Fast Real-Time PCR System (Applied Biosystems) using 5 µL of cDNA at 1 ng µL^-1^, 10 µL of the 2X PerfeCTa SYBR Green FastMix (Quantabio), 1 µL of primer pairs at 10 µM (final concentration of 250 nM for each primer) and 4 µL of nuclease-free water for a total volume of 20 µL. All samples were run in triplicate on a single 96-well plate for a given gene. We included no-template and no-reverse transcription controls. The thermocycling protocol was 95°C for 3 min (initial denaturation), followed by 40 cycles of 15 s at 95°C (denaturation) and 45 s at 60°C (annealing).

We used the NormFinder software (Andersen et al., 2004) to identify the optimal reference gene (or combination of reference genes) for our experiment between *ubc, hprt1, rpl13a, gapdh* and *β-actin* (Table 3). We did the analysis on 12 cDNA samples that were previously obtained in the same conditions as experimental samples during a pilot study. The NormFinder algorithm identified *ubc* as the most stable gene between time points. We thus calculated the relative expression of target genes using the **ΔΔ**Cq method adjusted for efficiency of each primer pairs (Pfaffl, 2001) with *ubc* as the reference gene.

**Table 3.**
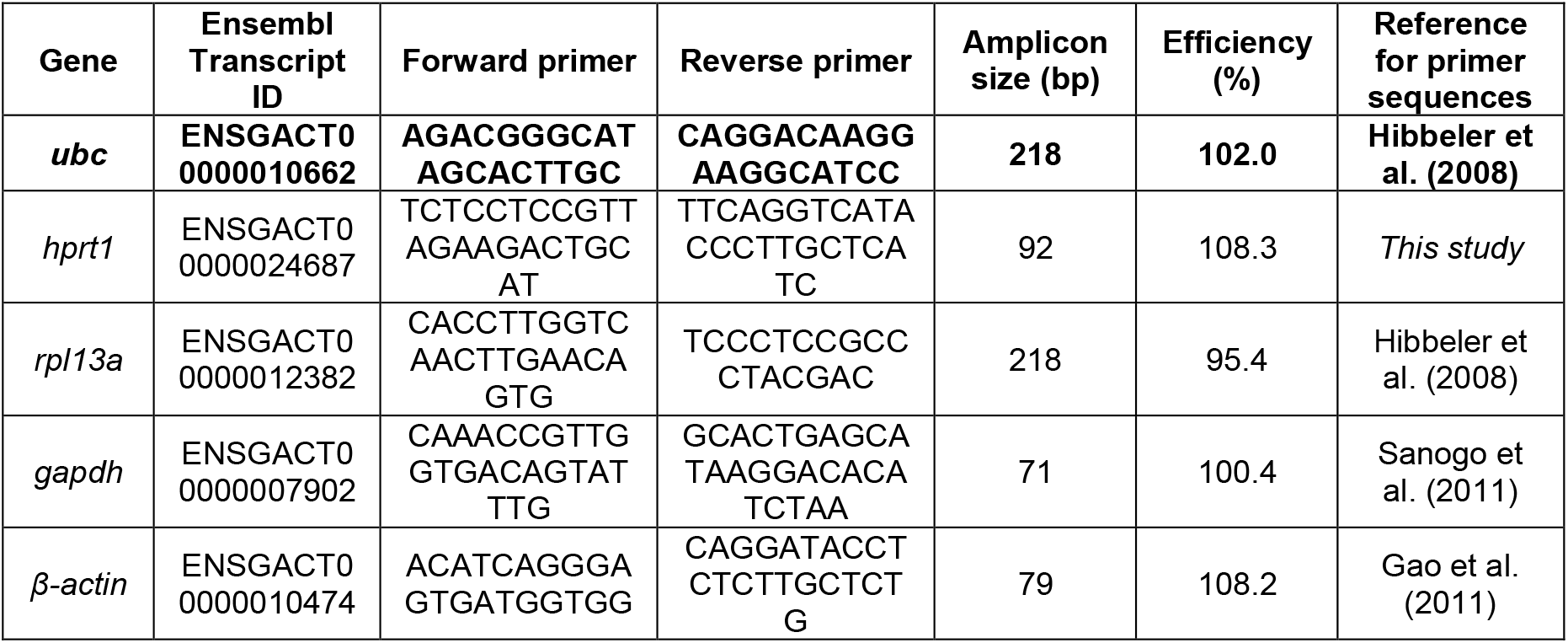
Characteristics of the primers used in the search for the optimal reference gene. Using the NormFinder software, *ubc* was identified as the most stable gene between time points and was used as the reference gene (shown in bold). Efficiency (E) was calculated using E = (10 ^-1/slope^ −1)*100 (Pfaffl, 2001).

### Data analysis

#### Locomotor activity rhythm

Of the 54 individuals that we used in our experiments, six were discarded from analyses because they died during experiments (n=3) or were parasitized (n=3). For the 48 remaining individuals, we gathered locomotor activity data in 10-min bins for analysis purposes. Actograms, activity profiles and 𝒳^2^ periodograms were produced using the ActogramJ plugin in ImageJ (Schmid et al., 2011) for each fish under LD (n=48) and under DD (n=17). The 𝒳^2^ periodogram analysis calculates Qp values for multiple periods within a fixed range. The period with the highest Qp value corresponds to the estimated period of the rhythm. Since Qp has a probability distribution of 𝒳^2^ (with a P-1 degree of freedom, where P is the period), we can determine if the Qp value for the estimated period is significant with α=0.05 (Sokolove and Bushell, 1978). In other words, the periodogram analysis lets us know if the rhythm is significant and, if so, what is the period of this rhythm. We first did the periodogram analysis using periods ranging between 0 h and 32 h, but we did not find any significant ultradian endogenous rhythms (i.e. rhythms with periods shorter than circadian rhythms). Thus, we show periodograms with periods ranging between 16 h and 32 h to facilitate visualization. We performed all other statistical analyses using R software version 4.0.1 (R Core Team, 2020). When needed, we evaluated normal distribution of data using Q-Q plots and Shapiro-Wilk test and we verified homogeneity of variances using Levene’s test.

#### Masking effect of light

We evaluated the difference in the average activity level (movements/10 min) during the light phase in LD and the subjective light phase in DD using a paired t-test. This comparison allows us to assess the importance of the masking effect of light, which can suppress or enhance activity without entraining the internal clock (Mrosovsky, 1999). We also verified if the difference in the average activity level during the light phase in LD and the subjective light phase in DD differs between sexes using a t-test.

#### Phase of activity

Although our hypothesis was that sticklebacks are diurnal, we rather observed a tendency towards nocturnality under LD. Thus, to quantify the phase of activity in each fish, we calculated the percentage of the total daily activity performed during the dark phase (also referred as the night activity). By assessing the night activity, we observed large interindividual variation for the phase of activity, but we also noticed large interindividual variation in total daily activity. We thus assessed sex differences in night activity and in total daily activity using t-tests and we evaluated the correlation between these two variables using Pearson’s correlation test. Data are represented as mean ± standard error of the mean.

#### Clock gene expression rhythms in the brain

Among the six individuals that were discarded from the analysis, five were from groups 2 and 3, so there were 31 individuals left for the brain gene expression analysis. We thus sampled eight individuals at 6:00, 18:00 and 0:00 (n=8) and seven individuals at 12:00 (n=7). Moreover, one individual was removed from the 18:00 time point for *clock2* because it was identified as an extreme outlier using the identify_outliers() function from the rstatix package in R (Kassambara, 2020). We evaluated differences in relative gene expression between time points using one-way ANOVA. Relative gene expression was also subjected to cosinor analysis using the cosinor2 package (Mutak, 2018). The cosinor analysis fits a cosine function with a known period (24 h) to the expression values so that we can estimate the amplitude, the acrophase (peak time) and the mesor (mean of all expression values) of the rhythm (Refinetti et al., 2007). This procedure also calculates the probability that the amplitude is significantly different from zero using the *F*-distribution. When the p-value is <0.05, we can consider that gene expression shows significant circadian rhythmicity.

## RESULTS

### Locomotor activity rhythm

Under a 12 h light:12 h dark cycle (LD), a significant daily rhythmicity of 24.0 h (𝒳^2^ periodogram analysis, p<0.05) was observed for all fish (Fig. 2). Under constant darkness (DD), most individuals were arrhythmic (Fig. 3A, C, E) and only three out of seventeen sticklebacks (18%) showed significant circadian rhythmicity (𝒳^2^ periodogram analysis, p<0.05, Fig. 3B, D, F) with periods of 24.8 h, 25.0 h and 26.3 h.

**Fig. 2.**
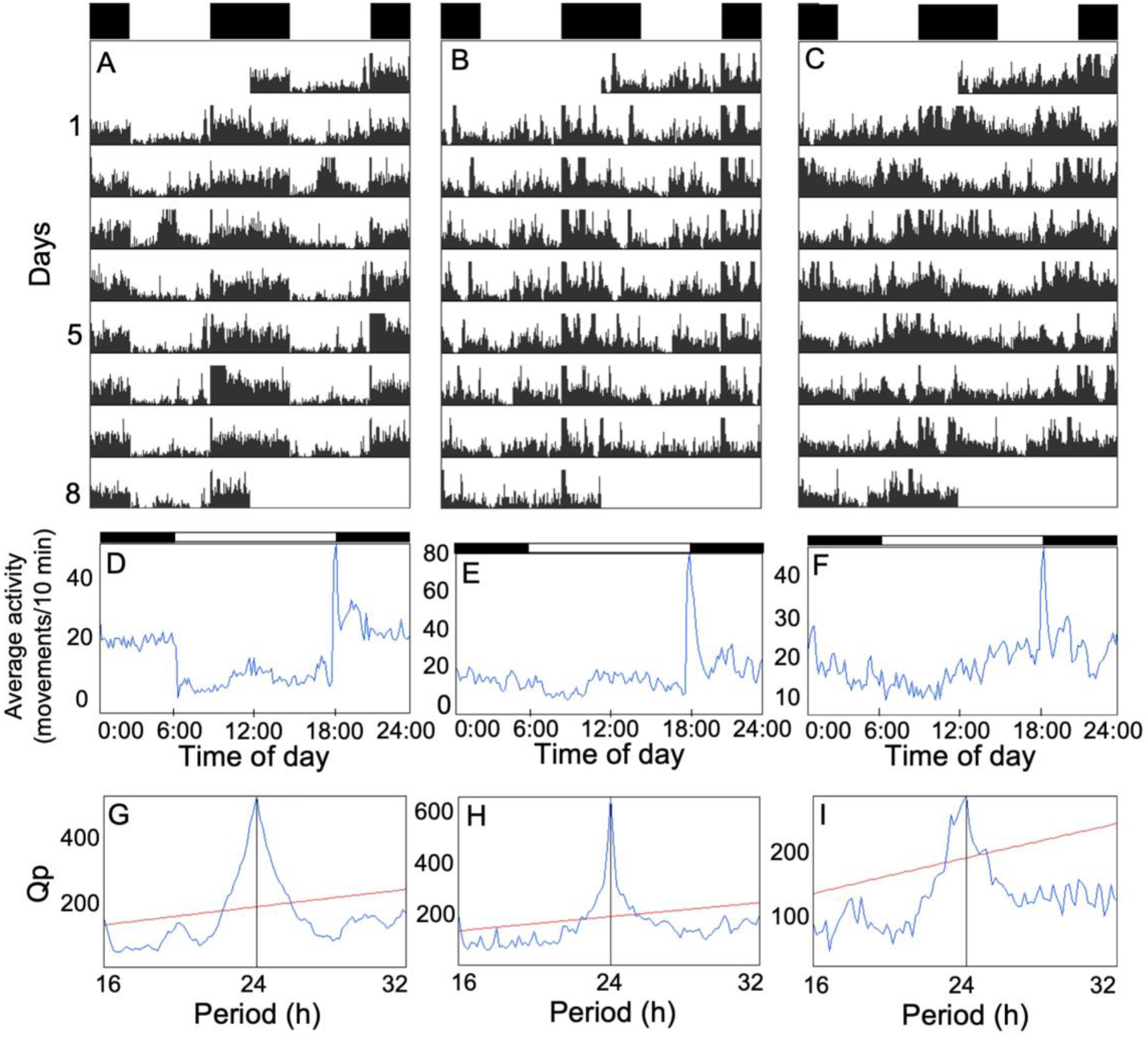
Under LD, sticklebacks display significant daily rhythmicity, but show variable activity patterns. Double-plotted actograms (A-C) of three representative individuals under a 12h light:12h dark cycle (LD) and their corresponding activity profile (D-F) and 𝒳^2^ periodogram (G-I). The white and black bars at the top of the actograms and the activity profiles represent the light and dark phases, respectively. From left to right, individuals display respectively 77%, 65% and 55% of their daily activity during the dark phase. Activity profiles show the average locomotor activity (number of movements) for each 10-min bin over the 8 days in LD. Qp values on the 𝒳^2^ periodograms quantify the rhythmic component of the activity and the red horizontal line indicates the significance threshold (set at p=0.05).

**Fig. 3.**
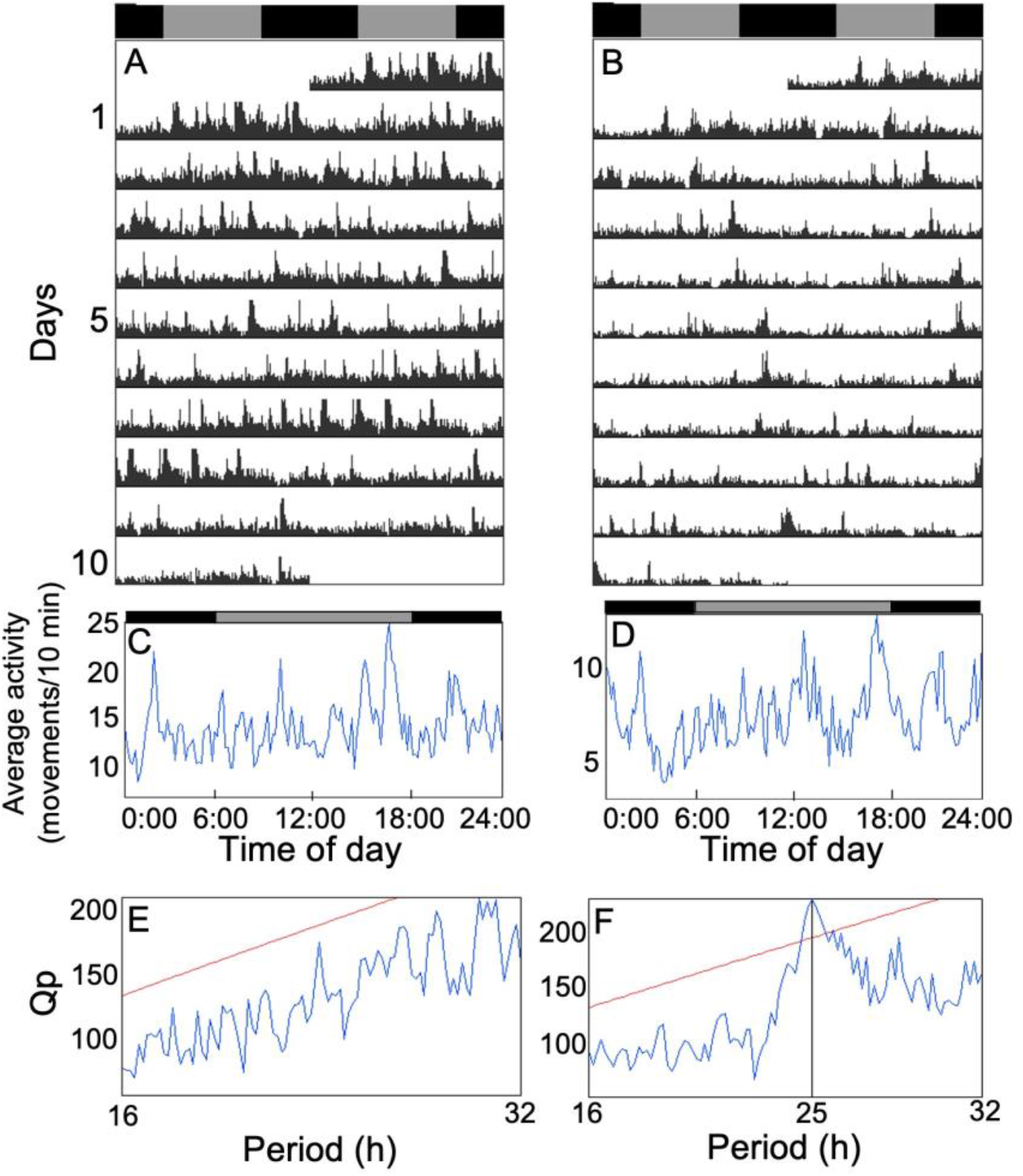
Under DD, most individuals are arrhythmic and only a few individuals show circadian rhythmicity. Double-plotted actograms (A-B) of two representative individuals under constant darkness (DD) and their corresponding activity profile (C-D) and 𝒳^2^ periodogram (E-F). The gray and black bars at the top of the actograms and activity profiles represent the subjective light and dark phases, respectively. Under DD, most sticklebacks do not display circadian rhythmicity, as represented by the individual on the left of the figure. On the right, we show one of the three individuals who exhibit significant circadian rhythmicity. Activity profiles show the average locomotor activity (number of movements) for each 10-min bin over the 10 days. Qp values on the 𝒳^2^ periodograms quantify the rhythmic component of the activity and the red horizontal line indicates the significance threshold (set at p=0.05).

### Masking effect of light

The average activity level (movements/10 min) was significantly lower during the light phase in LD than during the subjective light phase in DD (paired t-test, p<0.001, n=17, Fig. S3). The difference in the average activity level during the light phase in LD and the subjective light phase in DD was not significantly different between males (n=7) and females (n=10) (t-test, p=0.3).

### Phase of activity

Under LD, a few sticklebacks showed a well-defined phase of activity and were almost strictly nocturnal (Fig. 2A, D). However, most individuals displayed an unclear phase of activity and were just slightly more active during the night than during the day (Fig. 2B, C, E, F). On average, sticklebacks displayed 61.8±1.3% (n=48) of their daily activity during the dark phase. There was interindividual variation in the phase of activity, as measured by the percentage of the total daily activity displayed during the dark phase (also referred as the night activity, Fig. 4A), with individuals spending 46.5% to 87.5% of their active time at night. Of note, the three fish that were rhythmic in DD (described above) were not among the most nocturnal fish, as they displayed on average 53.0%, 52.9% and 57.0% of their daily activity during the night under LD. There was no significant difference between males (n=22) and females (n=26) in night activity (t-test, p=0.3). Under LD, sticklebacks also showed large interindividual variation in the total daily activity ranging from around 550 to 2750 movements/day (Fig. 4B). Males (1655±99 movements/day, n=22) were significantly more active than females (1357±95 movements/day, n=26) (t-test, p=0.04). There was also a significant negative correlation between night activity and total daily activity (Pearson’s correlation test, r=-0.3, p=0.04, n=48) so that the most nocturnal fish were also the least active (Fig. 4C).

**Fig. 4.**
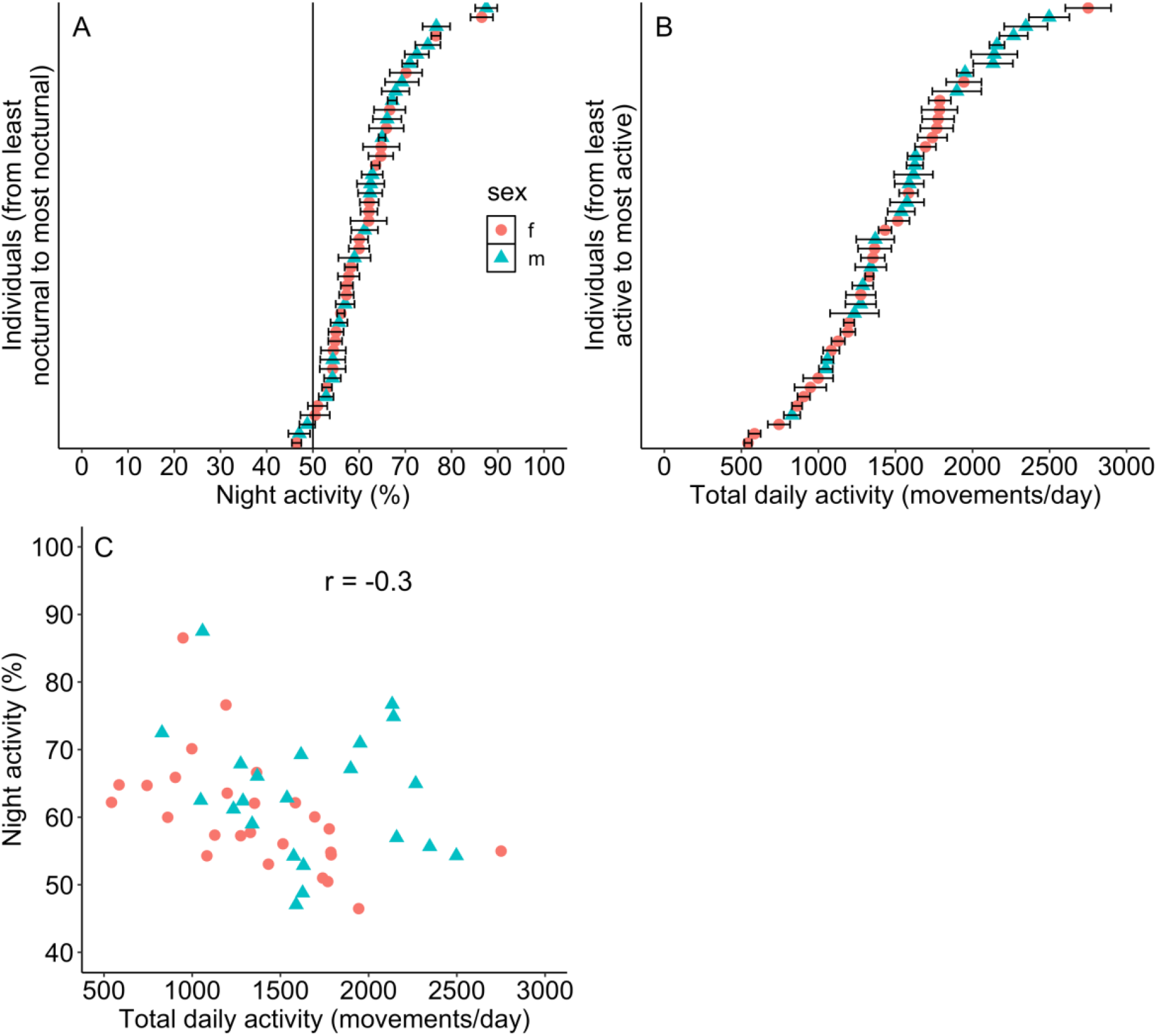
Under LD, sticklebacks are mostly nocturnal, but show large interindividual variation in the phase of activity and in the total daily activity. The most nocturnal fish are also the least active. (A) Average night activity of each individual under a 12 h light:12 h dark cycle (LD) for 8 days. Night activity corresponds to the percentage of the total daily activity displayed during the dark phase. Error bars represent the standard error of the mean. (B) Average total daily activity (movements/day) of each individual under LD for 8 days. (C) Correlation between the night activity (%) and the total daily activity (movements/day) under LD (average for 8 days). Note that axes are not starting at zero. Pearson’s correlation test, r=-0.3, p=0.04, n=48.

### Clock gene expression rhythms in the brain

We did not find any significant time-of-day variation in the relative expression of *bmal1a, clock1b, clock2, per1b* and *cry1b* in the brain of sticklebacks (one-way ANOVA, p>0.05, Fig. 5). In addition, the cosinor analysis did not detect any significant circadian rhythmicity in the relative expression of the five genes (cosinor analysis, p>0.05).

**Fig. 5.**
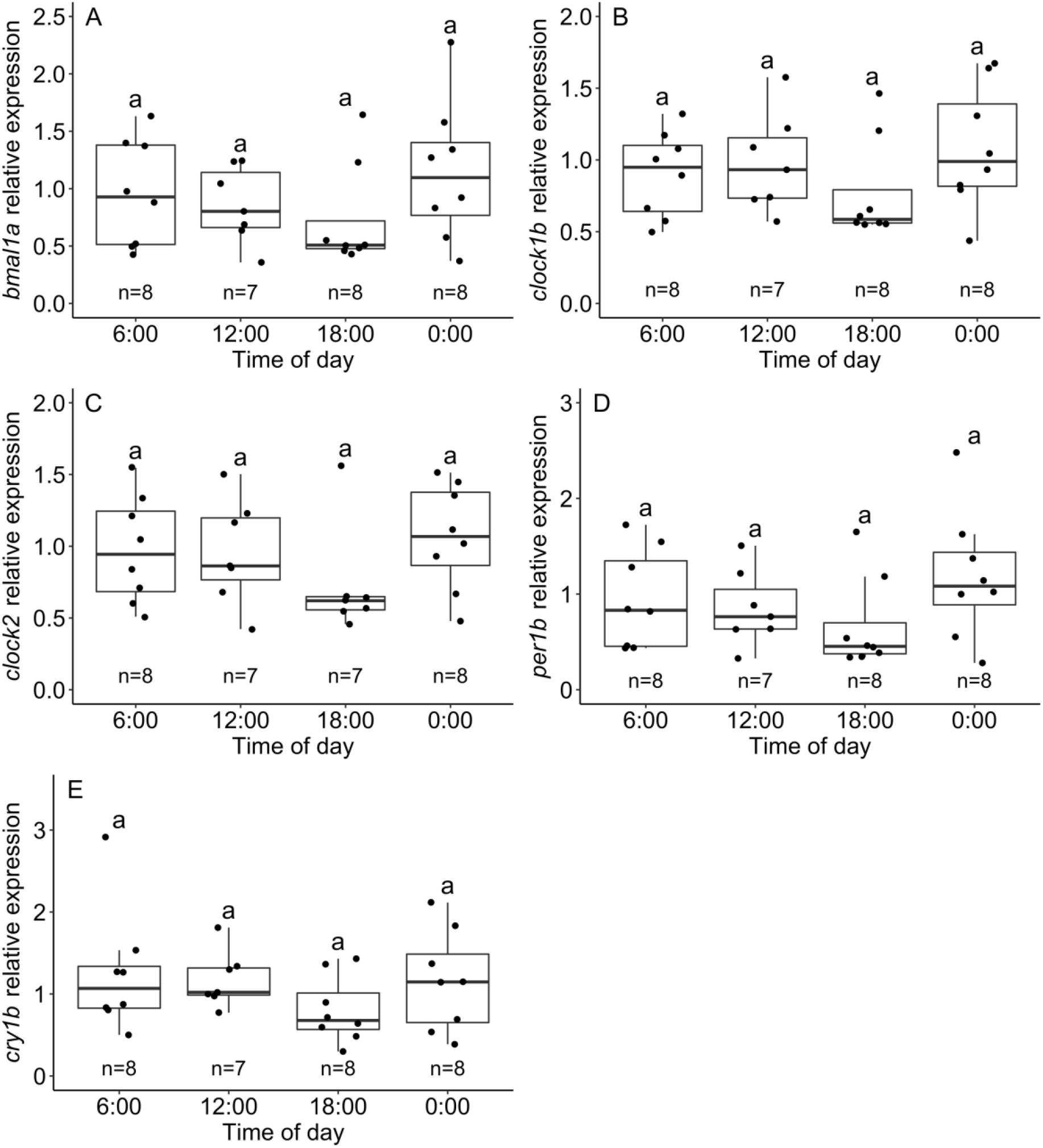
The expression of clock genes does not significantly vary during a 24-hour period in the brain of sticklebacks in DD. Time-of-day-dependent relative expression of *bmal1a* (A), *clock1b* (B), *clock2* (C), *per1b* (D) and *cry1b* (E) measured by qPCR in the brain of sticklebacks after one day in constant darkness (DD). The letter “a” denotes the absence of significant difference between time points for the five genes (one-way ANOVA, p>0.05). The black line in the middle of each boxplot indicates the median and each dot represents an individual. Sample size is shown for each time point.

## DISCUSSION

The circadian clock is an internal timekeeping system shared by almost all living organisms and has been mostly studied in model organisms. While knowledge about the functional importance and the evolution of circadian rhythms in natural environments is mounting, combining studies at the behavioural and molecular levels in individuals from natural populations but in controlled experimental settings is still in its early phase. In this study, using wild-caught sticklebacks, we investigated the circadian clock of this species at the behavioural and molecular levels. Our first objective was to determine if the daily rhythm of locomotor activity is under circadian clock control using a manipulation of the photoperiod. Under LD, all fish exhibited significant daily rhythmicity, while under DD, only a few individuals remained rhythmic. This result indicates that the circadian clock controls the locomotor activity rhythm in only a few sticklebacks, revealing a noteworthy interindividual variation. Our second objective was to determine the phase of activity of sticklebacks under LD. Contrary to our hypothesis, sticklebacks were mostly nocturnal. However, we observed again large interindividual variation: some fish were almost exclusively nocturnal while others were just slightly more active during the night than during the day. This variation was negatively correlated with the total daily activity, meaning that the most nocturnal fish were also the least active. This result suggests that light suppresses activity more strongly in some individuals, making them the most nocturnal fish. Our third objective was to describe the molecular oscillation of five clock genes (*bmal1a, clock1b, clock2, per1b* and *cry1b*) in the brain of sticklebacks under DD. Contrary to our hypothesis, we reported a lack of circadian rhythmicity for the five genes in the brain, which could either indicate that clock gene expression is not endogenously controlled, or that there was a significant oscillation but that we were unable to detect it, as a result of the large biological variation observed among individuals or because of technical issues.

### Locomotor activity rhythm under constant darkness

We found striking interindividual variation in circadian rhythms of activity in threespine sticklebacks. Our finding that not all individuals display a significant circadian rhythm of locomotor activity has been reported previously in other fish species. For instance, under constant conditions (constant darkness or light), the percentage of rhythmic individuals was 57% in goldfish (*Carassius auratus*, Iigo and Tabata, 1996), 50% in Nile tilapia (*Oreochromis niloticus*, Vera et al., 2009), 42% in tench (*Tinca tinca*, Herrero et al., 2003) and 30% in sharpsnout seabream (*Diplodus puntazzo*, Vera et al., 2006). In our experiment, 18% of sticklebacks were rhythmic in DD. Thus, a lack of circadian control on the locomotor activity rhythm seems common in fish. An advantage of not being under the strict control of the circadian clock could be that it allows the fish to rapidly adjust their phase of activity when critical changes occur in the environment, such as a shift in food availability, predation risk, mating opportunities, presence of parasites, etc. This is demonstrated by jet lag in animals that are strongly influenced by their internal clock, such as humans: it takes several days to adjust the phase of activity to a new environment and this re-entrainment is associated with many negative effects on health and cognitive performance (Waterhouse et al., 2007). Thus, in fish populations facing a particularly fluctuating environment, the individuals may benefit from being flexible and able to adjust their phase of activity, rather than their activity being rigidly controlled by their internal timekeeping system. For instance, the stickleback population in lac Témiscouata has to cope with several aquatic and avian predators (Reimchen, 1994; Tessier et al., 2008). All these fish and birds likely forage at various moments during the day and might themselves change their phase of activity according to various environmental factors or throughout the year. Sticklebacks thus probably must deal with many conflictual – and sometimes unpredictable – daily patterns in predation risk. Indeed, the lac Témiscouata population shows strong anti-predator morphology and behaviour, even when laboratory-reared (Lacasse and Aubin-Horth, 2012). Having a flexible daily schedule could further help sticklebacks to deal with several types of predators. On the other hand, the fact that some individuals kept an activity rhythm in constant darkness highlights that the extensive interindividual variation seen in many traits in sticklebacks, such as personality (Huntingford, 1976; Bell, 2005; Aubin-Horth et al., 2012), is also present in their circadian rhythms. Whether the variation quantified in these wild individuals arises from genetic variation or developmental plasticity in their early environment will need to be tested using common-environment experiments (Greenwood et al., 2011; Di-Poi et al., 2014). This interindividual variation suggests the hypothesis that there is more than one successful way to regulate its daily activities in that environment.

For the majority of the individuals that were not rhythmic in constant darkness, a lack of circadian regulation does not mean that they do not have a functional clock. It is possible that the clock molecular oscillator is partially uncoupled from the effectors, e.g. the locomotor system. For instance, uncoupling between clock gene expression rhythm and behavioural rhythm have been reported in the Mexican blind cavefish (*Astyanax mexicanus*, Beale et al., 2013). Similarly, the neuronal activity in the suprachiasmatic nucleus (the clock master oscillator in mammals) of guinea pigs (*Cavia porcellus*) shows robust circadian rhythmicity, but the animals express very unclear and weak activity rhythm (Kurumiya and Kawamura, 1988). It is thus possible that daily activities are not regulated by the clock molecular oscillator in sticklebacks as well. If this is the case, other behaviours, or physiological processes – such as the daily variation in the melatonin level – would be expected to be controlled by the circadian clock. It is also possible that other environmental factors entrain the circadian clock of sticklebacks. For instance, food availability was shown to entrain circadian locomotor activity rhythms in goldfish (*Carassius auratus*, Sánchez-Vázquez et al., 1997), tench (*Tinca tinca*, Herrero et al., 2005) and zebrafish (*Danio rerio*, Lopez-Olmeda et al., 2010). In our study, sticklebacks could only be entrained by the light-dark cycle since they were fed at random time and all other environmental cues were held constant. In future studies, asking whether other environmental factors can entrain circadian rhythms in sticklebacks would help us to understand what temporal cues are important for these fish in their natural environment. Alternatively, the photoperiodic conditions we used might have been inadequate and it could have led us to mistakenly think that the light-dark cycle could entrain the circadian clock in only a few sticklebacks. For instance, under LD, transitions between the light and the dark phases were very sudden, which is obviously not the case in nature since the sun sets and rises progressively. The sharp increase in activity observed every day just after the lights were turned off might indicate that this event was stressful for the fish. In future experiments, using a light gradient at sunrise and at sunset could help to better reproduce natural conditions (ex.: Lazado et al., 2014).

### Masking effect of light and phase of activity under light-dark cycle

Having established that the locomotor activity rhythm of sticklebacks is not controlled by the internal clock in most individuals, our results suggest that the masking effect of light contributes to the significant daily rhythm that we observed for all fish under LD. The masking effect of light refers to the direct influence of the photic signal on an organism’s behaviour, that is to say without the entrainment of its internal clock (Mrosovsky, 1999). As sticklebacks were generally more nocturnal under LD, the masking effect of light should suppress activity in this species (Mrosovsky, 1999). This is exactly what we observed: sticklebacks were less active during the light phase in LD than during the subjective light phase in DD (same hours of the day but different lighting conditions). This result indicates that light suppresses activity in sticklebacks, the definition of a masking effect.

We had hypothesized that sticklebacks are diurnal based on the fact that their visual opsins (Rennison et al., 2012) correspond to those of diurnal fish (Carleton et al., 2020) and that they are mostly captured during the day in the wild (Worgan and FitzGerald, 1981; Sjoberg, 1985; Reebs et al., 1995). However, we found that sticklebacks were, on average, mostly nocturnal under LD. The fact that some fish were almost strictly nocturnal suggests that sticklebacks can find food at night, either using visual or chemical cues (which has already been suggested by Mussen and Peeke, 2001). Whether sticklebacks do perform night activity in the wild is not known and might depend on several factors. In the laboratory, some sticklebacks might have chosen to be active during the night because they did not need to extensively rely on their visual system to find food (as their tank was quite small) and because they perceived the dark phase as safer. It has been previously shown in some species that there can be differences between the phase of daily activity in the laboratory and in the natural environment (reviewed in Calisi and Bentley, 2009). For instance, while mice (*Mus musculus*) are known for their nocturnal behaviour in the lab, they show variable phases of activity and are sometimes even exclusively diurnal when they are held in a semi-natural environment (Daan et al., 2011). Therefore, it is possible that sticklebacks are nocturnal in the lab and diurnal in their natural environment, and this could be verified using acoustic telemetry (March et al., 2010; Hussey et al., 2015; Alós et al., 2017).

We reported large interindividual variation in the phase of activity under LD. Interindividual differences in the phase of activity have often been reported in fish (reviewed in Reebs, 2002). For example, under LD, some Nile tilapia (*Oreochromis niloticus*) are diurnal, others are nocturnal, and some are active around the clock (Vera et al., 2009). Similar behaviours have been reported in goldfish (*Carassius auratus*, Iigo and Tabata, 1996) and in Atlantic salmon (*Salmo salar*, Richardson and McCleave, 1974). In line with these results, we showed that some sticklebacks were clearly nocturnal under LD, while others displayed an unclear phase of activity and were just slightly more active during the night than during the day. Large interindividual variation in the phase of daily activity thus also seems common in fish. In this study, we also observed that sticklebacks who restricted their daily activity to the dark phase were also the least active. The masking effect of light could thus be involved: some fish were more nocturnal because light suppressed their activity more strongly than that of the other fish. Observation of less active individuals in wild populations has already been reported in other fish species (Slavík and Horký, 2012; Závorka et al., 2016; Alós et al., 2017). Moreover, in accordance with our results, it has been shown that the less active fish react more to variations in light intensity than the more active individuals in wild brown trout juveniles (*Salmo trutta*) (Závorka et al., 2016). The ultimate cause of this interindividual variation is not known, but it could be that some fish have less energy to invest in activity and need to optimize the timing of their daily activity. They would thus benefit from being strongly affected by the light signal because it would allow them to only be active at the most optimal time of the day, which seems to be during the night for sticklebacks in our experiment. We must also consider the fact that our activity measure may be affected by a technical issue. The photoelectric sensors used only covered a portion of the tanks. Thigmotaxis or “wall-hugging” is a stress-related behaviour found in fish as in mammals (Maximino et al., 2010). If some individuals were more anxious than others in our study, they might have swum very close to the wall of their tank and been less detected by the sensor. Moreover, if some fish perceived the light phase as riskier, their thigmotaxis behaviour might have been more pronounced during the day than during the night. Therefore, the fish that we detected as less active and more nocturnal might have been as active as the other fish and active around the clock like the other fish, the only difference being that they would have spent more time swimming close to the wall of their tank during the day. In future experiments, this bias could be avoided by using more than one photoelectric sensor on each tank. Under LD, we also observed a significant sex difference in the total daily activity: males were more active than females. We reported that light did not suppress activity differently between sexes, so the masking effect of light is not in cause. One potential explanation is that males invest more energy in their daily activities because they have a higher energetic demand (Chmura et al., 2020) and forage more than females to find food in their tank. Another potential explanation is that if males were less anxious than females in our study, our activity monitoring system might have detected them more (as explained above). Lower anxiety levels in males than in females have been reported, for instance, in humans (Donner and Lowry, 2013) and in fish (Fontana et al., 2020). In summary, our results suggest that circadian and daily locomotor activity rhythms display large interindividual variation in sticklebacks, which seems to be a common feature in fish (Reebs, 2002). As mammals tend to exhibit more robust circadian behavioural rhythms (although there are exceptions: Bloch et al., 2013; Hazlerigg and Tyler, 2019), our study highlights the importance of investigating a wide diversity of species to better understand the evolution of circadian clocks.

### Clock gene expression rhythms in the brain

We did not detect any significant circadian rhythmicity in the relative expression of core clock genes in the brain of sticklebacks under constant darkness (DD), which suggests that either the molecular oscillator is highly light-dependent or that there was a significant oscillation but we were unable to detect it. The first interpretation implies that clock gene expression rhythms are not endogenously controlled, which contrasts with what has been observed in the brain or neural tissues of many other fish species (Whitmore et al., 1998; Cermakian et al., 2000; Patiño et al., 2011; Vera et al., 2013; Moore and Whitmore, 2014; Costa et al., 2016; Ceinos et al., 2019). A more parsimonious explanation is that a biological or technical effect prevented us from detecting any significant rhythmicity. First, it is possible that sticklebacks displayed interindividual variation in their acrophases (i.e. different peak times) of clock gene expression, so that the variation at each time point was too great to allow detection of a significant rhythm. Interestingly, interindividual variation in peak times of clock gene expression is often reported in natural populations, for example in humans (Teboul et al., 2005; Nováková et al., 2013; Ferrante et al., 2015; Takahashi et al., 2018). In fish, clock gene expression has not been quantified often in wild-caught populations, with the notable exception of the Mexican cavefish (*Astyanax mexicanus*) (Beale et al., 2013). Without surprise, it was shown that a wild population of Mexican cavefish displayed greater interindividual variation in clock gene expression than a laboratory population, a result that could be explained by greater genetic variation in the wild population (Beale et al., 2013). To demonstrate that wild sticklebacks display different peak times of clock gene expression, the same fish would have to be sampled multiple times over a 24-h period. As the sampling would need not to be lethal, using fin samples could be considered (Cavallari et al., 2011; Beale et al., 2013; Mogi et al., 2017).

Technical issues could also explain the fact that we did not detect significant circadian rhythmicity. We quantified clock gene expression in the whole brain, but if different regions of the stickleback brain host independent molecular oscillators that display different circadian rhythms or if some brain tissues are arrhythmic, using the whole brain might have drowned the rhythmic signal. For instance, previous studies in mammals reported that the same clock gene can have various peak times of expression in different brain regions (Abe et al., 2002; Mure et al., 2018). In fish, few studies have quantified clock gene expression in different brain regions, the size of this organ often being limiting. Among those who did, some reported distinct expression peaks between brain regions (Cermakian et al., 2000; Huang et al., 2010), but several others rather observed similar expression peaks throughout the brain (Whitmore et al., 1998; Weger et al., 2013; Moore and Whitmore, 2014; Costa et al., 2016). Besides, whole brains have often been used successfully to quantify clock gene expression rhythms in fish, both under LD (Lopez-Olmeda et al., 2010; Sánchez et al., 2010; Wang et al., 2015; Tudorache et al., 2018) and under DD (Whitmore et al., 1998; Cermakian et al., 2000; Vera et al., 2013; Moore and Whitmore, 2014). It thus seems that we could have detected significant rhythmicity using the whole brain of sticklebacks. That being said, in future studies, it would be possible to sample specific regions of the stickleback brain such as the diencephalon (which contains the hypothalamus) and the midbrain (which contains the optic tectum) (Sanogo et al., 2012; Greenwood and Peichel, 2015; Bell et al., 2016). These regions have been used a few times to quantify clock gene expression rhythms in other fish species (Feliciano et al., 2011; Martín-Robles et al., 2012; Moore and Whitmore, 2014; Costa et al., 2016). Another possibility would be to sample other organs such as the heart and the liver, which are commonly used to study the clock molecular oscillator in fish (Sánchez et al., 2010; Cavallari et al., 2011; Wang et al., 2015).

In this study, we showed that there is interindividual variation in the circadian rhythm of locomotor activity in wild sticklebacks, with most individuals exhibiting activity not controlled by their clock. In addition, we found that sticklebacks were mostly nocturnal under LD, but we observed large interindividual variation that could be due to a differential response to the masking effect of light among individuals. In future studies, asking whether a lack of circadian control is common in wild populations of sticklebacks or if it is driven by specific environmental challenges (such as high predation risk) will allow to better understand what selection pressures can shape the evolution of circadian rhythms. Moreover, assessing whether other biological rhythms are more strongly controlled by the clock and if the stickleback circadian system can be entrained by other environmental factors (such as food availability) will inform us about the functional importance of the circadian clock in this species. In parallel, studying the molecular oscillator will tell us what clock mechanisms underlie potential differences in circadian rhythms between populations and individuals. Importantly, in the study of gene expression, interindividual variation will need to be addressed and the choice of target organs used to quantify clock gene expression will affect the capacity to detect significant rhythmicity. Overall, our study suggests that a strong circadian control on behavioural rhythms is not necessarily advantageous in a natural population of sticklebacks and that the masking effect of light is potentially responsible for the large interindividual variation in the daily phase of activity.

## Acknowledgments

The authors would like to thank Dany Turcotte from Westburne Canada for his valuable advice regarding the photoelectric sensors and for graciously volunteering his time to program the controller. We thank Verônica A. Alves and Morgane Philippe for the field work, Sann Delaive for its help with tissue sampling and the personnel of the LARSEM at Université Laval for their help with fish rearing. We thank Chloé S. Berger, Florent Sylvestre and Verônica A. Alves for their comments on previous versions of this manuscript. We also thank Nicolas Cermakian for his valuable advice on this research.

## Supplementary information

**Fig. S1.**
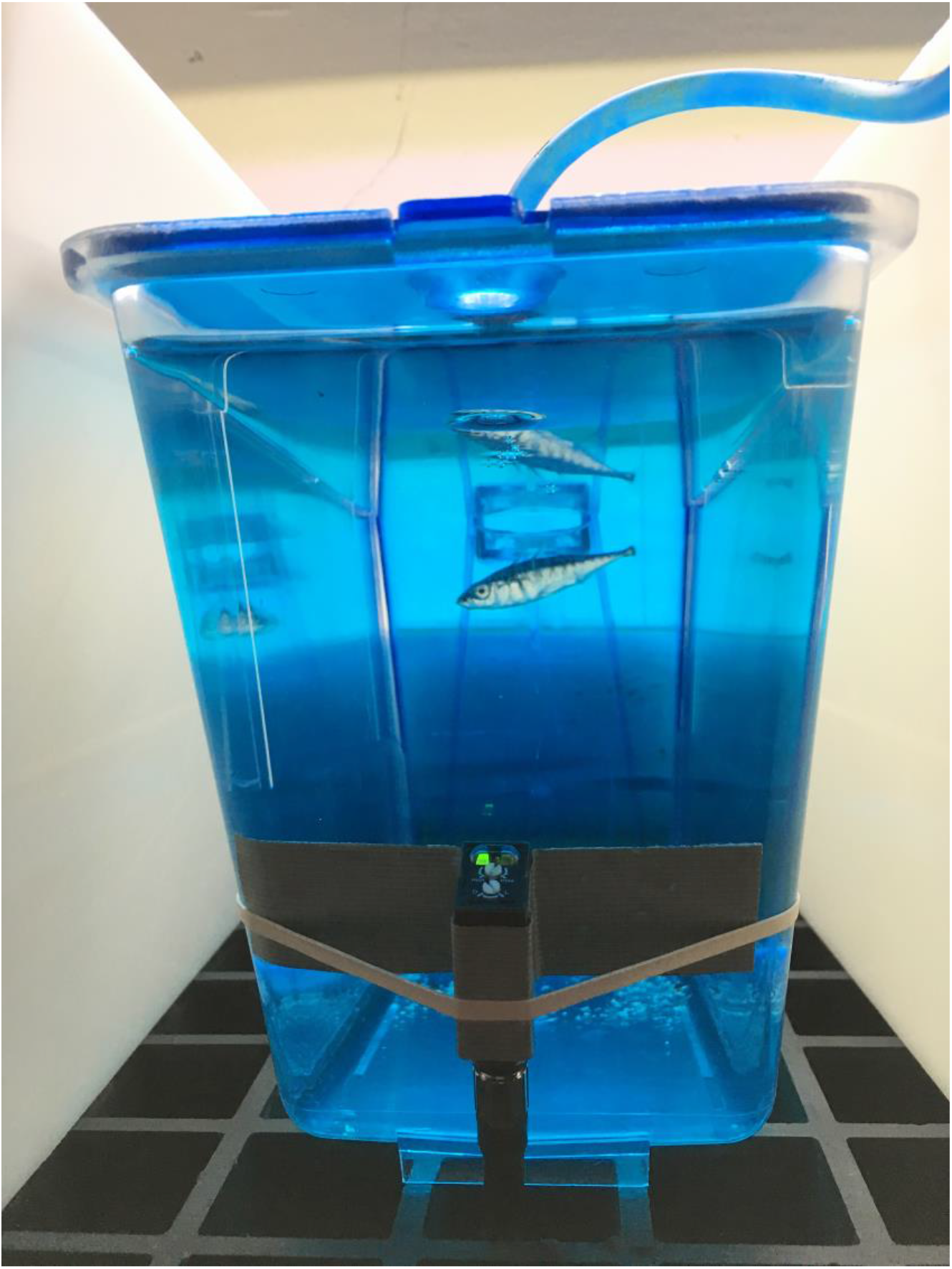
Position of the infrared photoelectric sensor on an experimental tank. Each experimental tank was equipped with one infrared photoelectric sensor placed in the lower third of the front wall. Every interruption of the infrared light beam by the fish was counted as one movement.

**Fig. S2.**
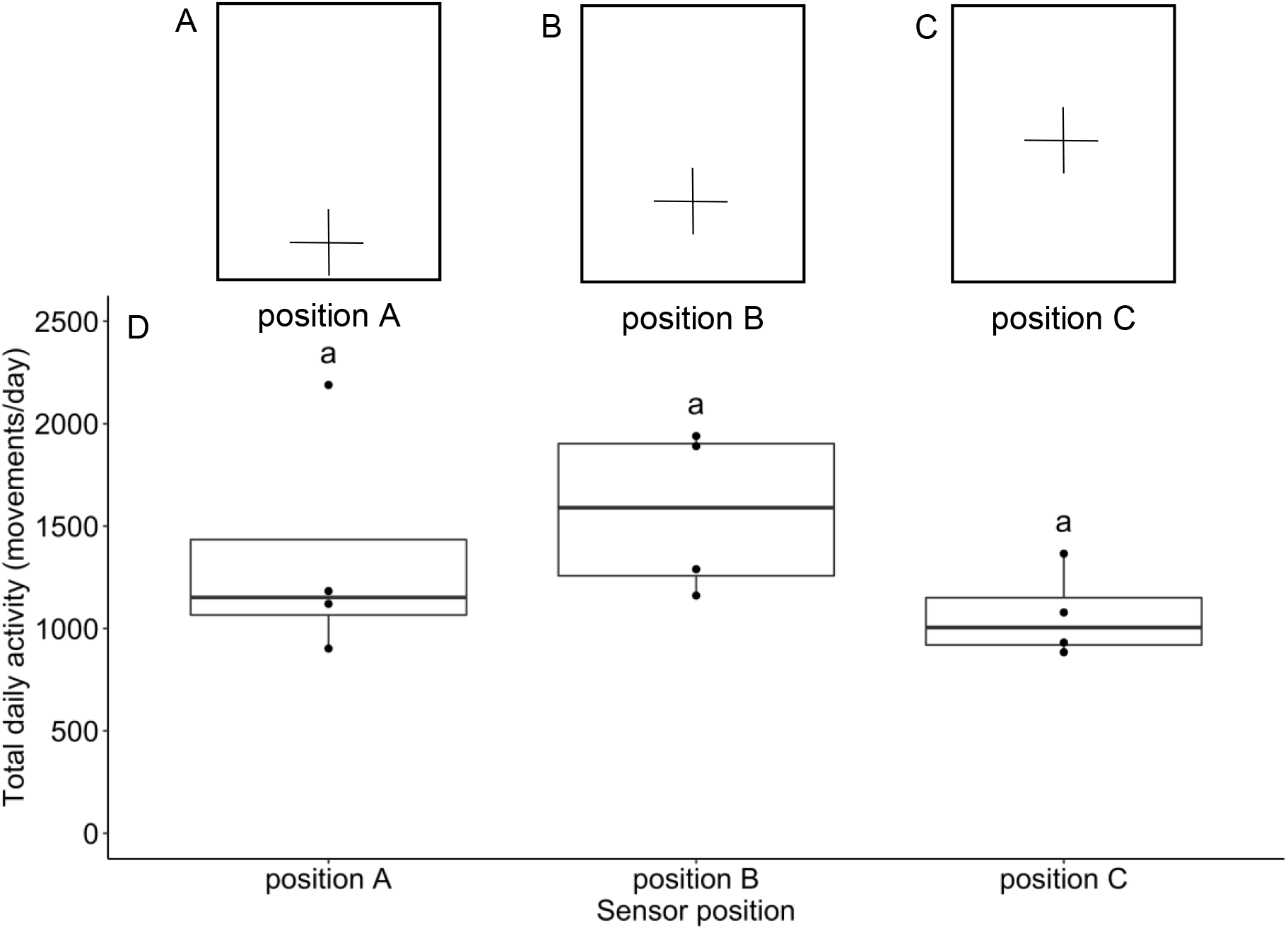
Sensors position optimization. (A, B, C) In order to determine in which position the sensors detect the most movements, we did a pilot study using 12 fish. We put each fish in an experimental tank equipped with a sensor that was either placed at the very bottom (position A), in the lower third (position B) or in the middle of the front wall (position C), so that there were 4 fish per position (n=4/position). We monitored locomotor activity for 8 days under a 12 h light:12 h dark cycle. The rectangle and the cross represent the front wall of the tank and the position of the sensor, respectively. (D) Average of the total daily activity (movements/day, average for 8 days) depending on the position of the sensor. Although there is no significant difference between positions (as indicated by the letter “a”, one-way ANOVA, p>0.05), the sensors in position B detect slightly more movements. We thus chose position B for our experiments (see Fig. S1). The black line in the middle of each boxplot indicates the median and each dot represents an individual.

**Fig. S3.**
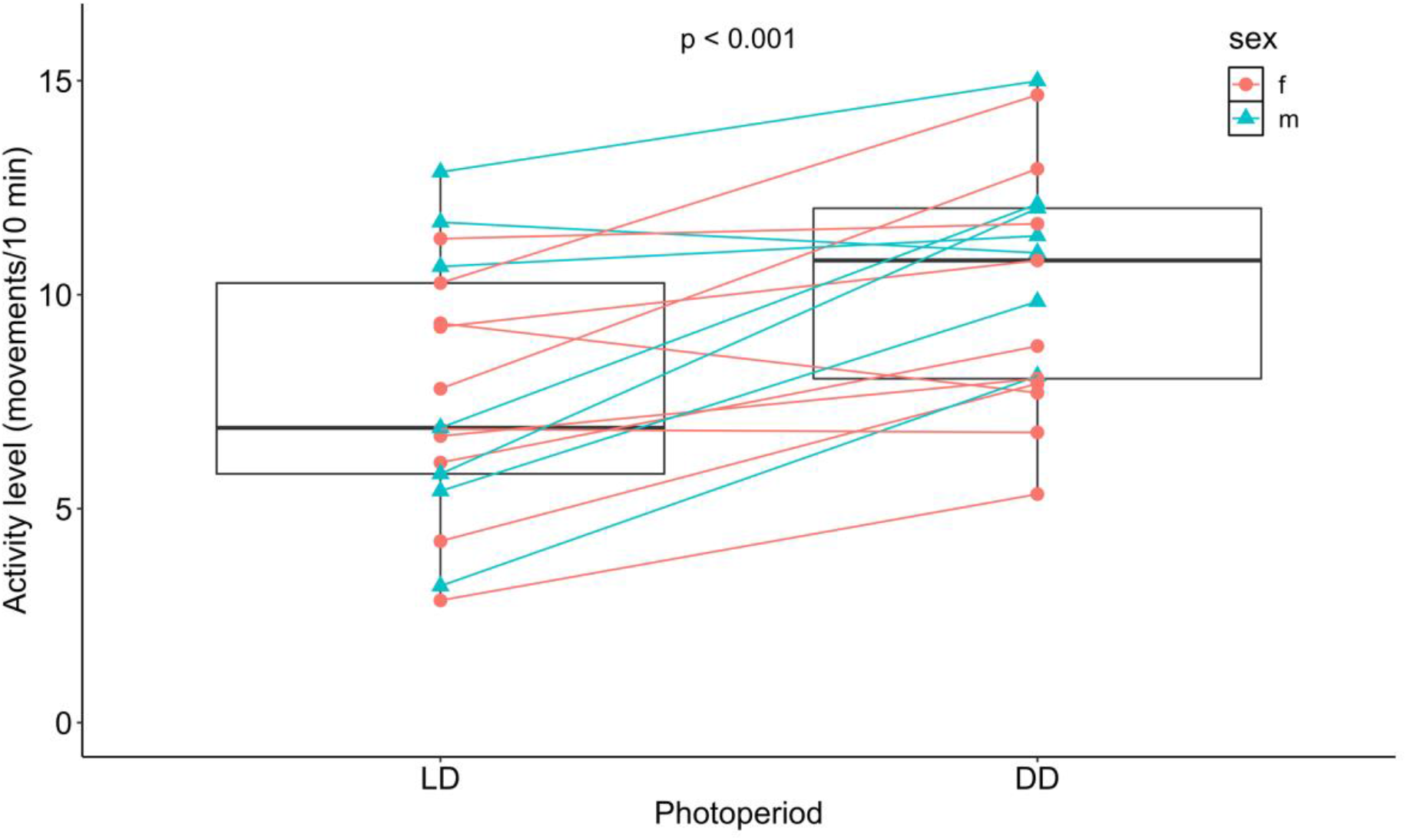
Sticklebacks are less active during the light phase in LD than during the subjective light phase in DD. Average activity level (movements/10 min) for each individual during the light phase of a 12 h light:12 h dark cycle (LD, average for 8 days) and during the subjective light phase in constant darkness (DD, average for 10 days). Paired t-test, p<0.001, n=17.

## References

Abe, M., Herzog, E. D., Yamazaki, S., Straume, M., Tei, H., Sakaki, Y., Menaker, M. and Block, G. D. (2002). Circadian rhythms in isolated brain regions. J. Neurosci. 22, 350–356.

Alós, J., Martorell-Barceló, M. and Campos-Candela, A. (2017). Repeatability of circadian behavioural variation revealed in free-ranging marine fish. R. Soc. open sci. 4, 160791.

Andersen, C. L., Jensen, J. L. and Ørntoft, T. F. (2004). Normalization of real-time quantitative reverse transcription-PCR data: A model-based variance estimation approach to identify genes suited for normalization, applied to bladder and colon cancer data sets. Cancer Res. 64, 5245–5250.

Aubin-Horth, N., Deschenes, M. and Cloutier, S. (2012). Natural variation in the molecular stress network correlates with a behavioural syndrome. Horm. Behav. 61, 140–146.

Bayarri, M. J., Muñoz-Cueto, J. A., López-Olmeda, J. F., Vera, L. M., Rol De Lama, M. A., Madrid, J. A. and Sánchez-Vázquez, F. J. (2004). Daily locomotor activity and melatonin rhythms in Senegal sole (Solea senegalensis). Physiol. Behav. 81, 577–583

Beale, A., Guibal, C., Tamai, T. K., Klotz, L., Cowen, S., Peyric, E., Reynoso, V. H., Yamamoto, Y. and Whitmore, D. (2013). Circadian rhythms in Mexican blind cavefish Astyanax mexicanus in the lab and in the field. Nat. Commun. 4, 2769.

Bell, A. M. (2005). Behavioural differences between individuals and two populations of stickleback (Gasterosteus aculeatus). J. Evol. Biol. 18, 464–473.

Bell, A. M., Bukhari, S. A. and Sanogo, Y. O. (2016). Natural variation in brain gene expression profiles of aggressive and nonaggressive individual sticklebacks. Behaviour 153, 1723–1743.

Bell, M. A. and Foster, S. A. (1994). The Evolutionary Biology of the Threespine Stickleback. Oxford, UK: Oxford University Press.

Bell-Pedersen, D., Cassone, V. M., Earnest, D. J., Golden, S. S., Hardin, P. E., Thomas, T. L. and Zoran, M. J. (2005). Circadian rhythms from multiple oscillators: lessons from diverse organisms. Nat. Rev. Genet. 6, 544–556.

Bloch, G., Barnes, B. M., Gerkema, M. P. and Helm, B. (2013). Animal activity around the clock with no overt circadian rhythms: patterns, mechanisms and adaptive value. Proc. R. Soc. B 280, 20130019.

Calisi, R. M. and Bentley, G. E. (2009). Lab and field experiments: Are they the same animal? Horm. Behav. 56, 1–10.

Carleton, K. L., Escobar-Camacho, D., Stieb, S. M., Cortesi, F. and Marshall, N. J. (2020). Seeing the rainbow: mechanisms underlying spectral sensitivity in teleost fishes. J. Exp. Biol. 223, jeb193334.

Cavallari, N., Frigato, E., Vallone, D., Frohlich, N., Lopez-Olmeda, J. F., Foa, A., Berti, R., Sánchez-Vázquez, F. J., Bertolucci, C. and Foulkes, N. S. (2011). A blind circadian clock in cavefish reveals that opsins mediate peripheral clock photoreception. PLoS Biol. 9, e1001142.

Ceinos, R. M., Chivite, M., López-Patiño, M. A., Naderi, F., Soengas, J. L., Foulkes, N. S. and Míguez, J. M. (2019). Differential circadian and light-driven rhythmicity of clock gene expression and behaviour in the turbot, Scophthalmus maximus. PLoS One 14, e0219153.

Cermakian, N., Whitmore, D., Foulkes, N. S. and Sassone-Corsi, P. (2000). Asynchronous oscillations of two zebrafish CLOCK partners reveal differential clock control and function. Proc. Natl. Acad. Sci. U. S. A. 97, 4339–4344.

Challet, E. (2007). Minireview: Entrainment of the suprachiasmatic clockwork in diurnal and nocturnal mammals. Endocrinology 148, 5648–5655.

Chiesa, J. J., Aguzzi, J., García, J. A., Sardà, F. and De La Iglesia, H. O. (2010). Light intensity determines temporal niche switching of behavioral activity in deep-water Nephrops norvegicus (Crustacea: Decapoda). J. Biol. Rhythms 25, 277–287.

Chmura, H. E., Zhang, V. Y., Wilbur, S. M., Barnes, B. M., Buck, C. L. and Williams, C. T. (2020). Plasticity and repeatability of activity patterns in free-living Arctic ground squirrels. Anim. Behav. 169, 81–91.

Costa, L. S., Serrano, I., Sánchez-Vázquez, F. J. and López-Olmeda, J. F. (2016). Circadian rhythms of clock gene expression in Nile tilapia (Oreochromis niloticus) central and peripheral tissues: influence of different lighting and feeding conditions. J. Comp. Physiol. B 186, 775–785.

Daan, S., Spoelstra, K., Albrecht, U., Schmutz, I., Daan, M., Daan, B., Rienks, F., Poletaeva, I., Dell’omo, G., Vyssotski, A., et al. (2011). Lab mice in the field: Unorthodox daily activity and effects of a dysfunctional circadian clock allele. J. Biol. Rhythms 26, 118–129.

Di Poi, C., Belanger, D., Amyot, M., Rogers, S. and Aubin-Horth, N. (2016). Receptors rather than signals change in expression in four physiological regulatory networks during evolutionary divergence in threespine stickleback. Mol. Ecol. 25, 3416–3427.

Di-Poi, C., Lacasse, J., Rogers, S. M. and Aubin-Horth, N. (2014). Extensive behavioural divergence following colonisation of the freshwater environment in threespine sticklebacks. PLoS One 9, e98980.

Dominoni, D. M., Åkesson, S., Klaassen, R., Spoelstra, K. and Bulla, M. (2017). Methods in field chronobiology. Philos. Trans. R. Soc. Lond. B Biol. Sci. 372, 20160247.

Donner, N. C. and Lowry, C. A. (2013). Sex differences in anxiety and emotional behavior. Pflugers Arch. 465, 601–626.

Falcón, J., Besseau, L., Fuentès, M., Sauzet, S., Magnanou, E. and Boeuf, G. (2009). Structural and functional evolution of the pineal melatonin system in vertebrates. Ann. N. Y. Acad. Sci. 1163, 101–111.

Feliciano, A., Vivas, Y., De Pedro, N., Delgado, M. J., Velarde, E. and Isorna, E. (2011). Feeding time synchronizes clock gene rhythmic expression in brain and liver of goldfish (Carassius auratus). J. Biol. Rhythms 26, 24–33.

Ferrante, A., Gellerman, D., Ay, A., Woods, K. P., Filipowicz, A. M., Jain, K., Bearden, N. and Ingram, K. K. (2015). Diurnal preference predicts phase differences in expression of human peripheral circadian clock genes. J. Circadian Rhythms 13, 1–7.

Fontana, B. D., Cleal, M. and Parker, M. O. (2020). Female adult zebrafish (Danio rerio) show higher levels of anxiety-like behavior than males, but do not differ in learning and memory capacity. Eur. J. Neurosci. 52, 2604–2613.

Fuchikawa, T., Eban-Rothschild, A., Nagari, M., Shemesh, Y. and Bloch, G. (2016). Potent social synchronization can override photic entrainment of circadian rhythms. Nat. Commun. 7, 11662.

Gao, K., Brandt, I., Goldstone, J. V. and Jönsson, M. E. (2011). Cytochrome P450 1A, 1B, and 1C mRNA induction patterns in three-spined stickleback exposed to a transient and a persistent inducer. Comp. Biochem. Physiol. C Toxicol. Pharmacol. 154, 42–55.

Greenwood, A. K., Jones, F. C., Chan, Y. F., Brady, S. D., Absher, D. M., Grimwood, J., Schmutz, J., Myers, R. M., Kingsley, D. M. and Peichel, C. L. (2011). The genetic basis of divergent pigment patterns in juvenile threespine sticklebacks. Heredity 107, 155–166.

Greenwood, A. K. and Peichel, C. L. (2015). Social regulation of gene expression in threespine sticklebacks. PLoS One 10, e0137726–e0137726.

Hanson, D., Hu, J., Hendry, A. P. and Barrett, R. D. H. (2017). Heritable gene expression differences between lake and stream stickleback include both parallel and antiparallel components. Heredity 119, 339–348.

Hazlerigg, D. G. and Tyler, N. J. C. (2019). Activity patterns in mammals: Circadian dominance challenged. PLoS Biol. 17, e3000360.

Helm, B., Visser, M. E., Schwartz, W., Kronfeld-Schor, N., Gerkema, M., Piersma, T. and Bloch, G. (2017). Two sides of a coin: ecological and chronobiological perspectives of timing in the wild. Philos. Trans. R. Soc. Lond. B Biol. Sci. 372, 20160246.

Herrero, M. J., Madrid, J. A. and Sánchez-Vázquez, F. J. (2003). Entrainment to light of circadian activity rhythms in tench (Tinca tinca). Chronobiol. Int. 20, 1001–17.

Herrero, M. J., Pascual, M., Madrid, J. A. and Sánchez-Vázquez, F. J. (2005). Demand-feeding rhythms and feeding-entrainment of locomotor activity rhythms in tench (Tinca tinca). Physiol. Behav. 84, 595–605.

Hibbeler, S., Scharsack, J. P. and Becker, S. (2008). Housekeeping genes for quantitative expression studies in the three-spined stickleback Gasterosteus aculeatus. BMC Mol. Biol. 9, 18.

Huang, T.-S., Ruoff, P. and Fjelldal, P. G. (2010). Effect of continuous light on daily levels of plasma melatonin and cortisol and expression of clock genes in pineal gland, brain, and liver in Atlantic salmon postsmolts. Chronobiol. Int. 27, 1715–1734.

Huntingford, F. A. (1976). The relationship between anti-predator behaviour and aggression among conspecifics in the three-spined stickleback, Gasterosteus Aculeatus. Anim. Behav. 24, 245–260.

Hussey, N. E., Kessel, S. T., Aarestrup, K., Cooke, S. J., Cowley, P. D., Fisk, A. T., Harcourt, R. G., Holland, K. N., Iverson, S. J., Kocik, J. F., et al. (2015). Aquatic animal telemetry: A panoramic window into the underwater world. Science 348, 1255642.

Hut, R. A., Kronfeld-Schor, N., Van Der Vinne, V. and De La Iglesia, H. (2012). In search of a temporal niche: Environmental factors. In Progress in Brain Research (ed. A. Kalsbeek, M. Merrow, T. Roenneberg and R. G. Foster), pp. 281–304. Oxford, UK: Elsevier.

Iigo, M. and Tabata, M. (1996). Circadian rhythms of locomotor activity in the goldfish Carassius auratus. Physiol. Behav. 60, 775–81.

Ishikawa, A., Kabeya, N., Ikeya, K., Kakioka, R., Cech, J. N., Osada, N., Leal, M. C., Inoue, J., Kume, M., Toyoda, A., et al. (2019). A key metabolic gene for recurrent freshwater colonization and radiation in fishes. Science 364, 886–889.

Kassambara, A. (2020). rstatix: Pipe-friendly framework for basic statistical tests. R package version 0.5.0.

Kitano, J., Lema, S. C., Luckenbach, J. A., Mori, S., Kawagishi, Y., Kusakabe, M., Swanson, P. and Peichel, C. L. (2010). Adaptive divergence in the thyroid hormone signaling pathway in the stickleback radiation. Curr. Biol. 20, 2124–2130.

Kronfeld-Schor, N., Bloch, G. and Schwartz, W. J. (2013). Animal clocks: when science meets nature. Proc. R. Soc. B 280, 20131354.

Kulczykowska, E., Kleszczyńska, A., Gozdowska, M. and Sokołowska, E. (2017). The time enzyme in melatonin biosynthesis in fish: Day/night expressions of three aralkylamine N-acetyltransferase genes in three-spined stickleback. Comp. Biochem. Physiol. A Mol. Integr. Physiol. 208, 46–53.

Kumar, V. (2017). Biological Timekeeping: Clocks, Rhythms and Behaviour. New Delhi, India: Springer India.

Kurumiya, S. and Kawamura, H. (1988). Circadian oscillation of the multiple unit activity in the guinea pig suprachiasmatic nucleus. J. Comp. Physiol. A 162, 301–308.

Lacasse, J. and Aubin-Horth, N. (2012). A test of the coupling of predator defense morphology and behavior variation in two threespine stickleback populations. Curr. Zool. 58, 53–65.

Lazado, C. C., Kumaratunga, H. P. S., Nagasawa, K., Babiak, I., Giannetto, A. and Fernandes, J. M. O. (2014). Daily rhythmicity of clock gene transcripts in atlantic cod fast skeletal muscle. PLoS One 9, e99172.

Liu, C., Hu, J., Qu, C., Wang, L., Huang, G., Niu, P., Zhong, Z., Hong, F., Wang, G., Postlethwait, J. H., et al. (2015). Molecular evolution and functional divergence of zebrafish (Danio rerio) cryptochrome genes. Sci. Rep. 5, 8113.

Lopez-Olmeda, J. F., Tartaglione, E. V., De La Iglesia, H. O. and Sánchez-Vázquez, F. J. (2010). Feeding entrainment of food-anticipatory activity and per1 expression in the brain and liver of zebrafish under different lighting and feeding conditions. Chronobiol. Int. 27, 1380–1400.

March, D., Palmer, M., Alós, J., Grau, A. and Cardona, F. (2010). Short-term residence, home range size and diel patterns of the painted comber Serranus scriba in a temperate marine reserve. Mar. Ecol. Prog. Ser. 400, 195–206.

Martín-Robles, Á. J., Whitmore, D., Sánchez-Vázquez, F. J., Pendón, C. and Muñoz-Cueto, J. A. (2012). Cloning, tissue expression pattern and daily rhythms of Period1, Period2, and Clock transcripts in the flatfish Senegalese sole, Solea senegalensis. J. Comp. Physiol. B 182, 673–685.

Maximino, C., De Brito, T. M., Da Silva Batista, A. W., Herculano, A. M., Morato, S. and Gouveia, A. (2010). Measuring anxiety in zebrafish: A critical review. Behav. Brain Res. 214, 157–171.

Mayer, I., Bornestaf, C., Wetterberg, L. and Borg, B. (1997). Melatonin does not prevent long photoperiod stimulation of secondary sexual characters in the male three-spined stickleback Gasterosteus aculeatus. Gen. Comp. Endocrinol. 108, 386–394.

Mckinnon, J. S., Kitano, J. and Aubin-Horth, N. (2019). Gasterosteus, Anolis, Mus, and more-the changing roles of vertebrate models in evolution and behaviour. Evol. Ecol. Res. 20, 1–25.

Mogi, M., Yokoi, H. and Suzuki, T. (2017). Analyses of the cellular clock gene expression in peripheral tissue, caudal fin, in the Japanese flounder, Paralichthys olivaceus. Gen. Comp. Endocrinol. 248, 97–105.

Moore, H. A. and Whitmore, D. (2014). Circadian rhythmicity and light sensitivity of the zebrafish brain. PLoS One 9, e86176.

Mrosovsky, N. (1999). Masking: history, definitions, and measurement. Chronobiol. Int. 16, 415–429.

Mure, L. S., Le, H. D., Benegiamo, G., Chang, M. W., Rios, L., Jillani, N., Ngotho, M., Kariuki, T., Dkhissi-Benyahya, O., Cooper, H. M., et al. (2018). Diurnal transcriptome atlas of a primate across major neural and peripheral tissues. Science 359, eaao0318.

Mussen, T. D. and Peeke, H. V. S. (2001). Nocturnal feeding in the marine threespine stickleback (Gasterosteus aculeatus L.): Modulation by chemical stimulation. Behaviour 138, 857–871.

Mutak, A. (2018). cosinor2: Extended tools for cosinor analysis of rhythms. R package version 0.2.1.

Nováková, M., Sládek, M. and Sumová, A. (2013). Human chronotype is determined in bodily cells under real-life conditions. Chronobiol. Int. 30, 607–617.

Ostlund-Nilsson, S., Mayer, I. and Huntingford, F. A. (2007). Biology of the Three-Spined Stickleback. Boca Raton, FL, US: CRC Press.

Pando, M. P., Pinchak, A. B., Cermakian, N. and Sassone-Corsi, P. (2001). A cell-based system that recapitulates the dynamic light-dependent regulation of the vertebrate clock. Proc. Natl. Acad. Sci. U. S. A. 98, 10178–10183.

Patiño, M. a. L., Rodríguez-Illamola, A., Conde-Sieira, M., Soengas, J. L. and Míguez, J. M. (2011). Daily rhythmic expression patterns of Clock1a, Bmal1, and Per1 genes in retina and hypothalamus of the rainbow trout, Oncorhynchus mykiss. Chronobiol. Int. 28, 381–389.

Peichel, C. L., Ross, J. A., Matson, C. K., Dickson, M., Grimwood, J., Schmutz, J., Myers, R. M., Mori, S., Schluter, D. and Kingsley, D. M. (2004). The master sex-determination locus in threespine sticklebacks is on a nascent Y chromosome. Curr. Biol. 14, 1416–1424.

Pellman, B. A., Kim, E., Reilly, M., Kashima, J., Motch, O., De La Iglesia, H. O. and Kim, J. J. (2015). Time-specific fear acts as a non-photic entraining stimulus of circadian rhythms in rats. Sci. Rep. 5, 14916.

Pfaffl, M. W. (2001). A new mathematical model for relative quantification in real-time RT-PCR. Nucleic Acids Res. 29, e45.

Pomianowski, K., Gozdowska, M., Burzyński, A., Kalamarz-Kubiak, H., Sokołowska, E., Kijewska, A. and Kulczykowska, E. (2020). A study of aanat and asmt expression in the three-spined stickleback eye and skin: Not only “on the way to melatonin”. Comp. Biochem. Physiol. A Mol. Integr. Physiol. 241, 110635.

Prokkola, J. M., Nikinmaa, M., Lubiana, P., Kanerva, M., Mccairns, R. J. S. and Gotting, M. (2015). Hypoxia and the pharmaceutical diclofenac influence the circadian responses of three-spined stickleback. Aquat. Toxicol. 158, 116–124.

Quinn, T. P., Sergeant, C. J., Beaudreau, A. H. and Beauchamp, D. A. (2012). Spatial and temporal patterns of vertical distribution for three planktivorous fishes in Lake Washington. Ecol. Freshw. Fish 21, 337–348.

R Core Team (2020). R: A language and environment for statistical computing. R Foundation for Statistical Computing.

Reebs, S. G. (2002). Plasticity of diel and circadian activity rhythms in fishes. Rev. Fish. Biol. Fish. 12, 349–371.

Reebs, S. G., Boudreau, L., Hardie, R. and Cunjak, R. A. (1995). Diel activity patterns of lake chubs and other fishes in a temperate stream. Can. J. Zool. 73, 1221–1227.

Reebs, S. G., Jr., F. G. W. and Fitzgerald, G. J. (1984). Diel patterns of fanning activity, egg respiration, and the nocturnal behavior of male three-spined sticklebacks, Gasterosteus aculeatus L. (f. trachurus). Can. J. Zool. 62, 329–334.

Refinetti, R. (2016). Daily and circadian rhythms. In Circadian Physiology (ed. R. Refinetti), pp. 169–236. Boca Raton: CRC Press.

Refinetti, R., Cornelissen, G. and Halberg, F. (2007). Procedures for numerical analysis of circadian rhythms. Biol. Rhythm Res. 38, 275–325.

Reimchen, T. E. (1994). Predators and morphological evolution in threespine stickleback. In The Evolutionary Biology of the Threespine Stickleback (ed. A. M. Bell and S. A. Foster), pp. 240–276. Oxford, UK: Oxford University Press.

Rennison, D. J., Owens, G. L. and Taylor, J. S. (2012). Opsin gene duplication and divergence in ray-finned fish. Mol. Phylogenet. Evol. 62, 986–1008.

Richardson, N. E. and Mccleave, J. D. (1974). Locomotor activity rhythms of juvenile atlantic salmon (Salmo salar) in various light conditions. Biol. Bull. 147, 422–32.

Rosensweig, C. and Green, C. B. (2020). Periodicity, repression, and the molecular architecture of the mammalian circadian clock. Eur. J. Neurosci. 51, 139–165.

Sánchez, J. A., Madrid, J. A. and Sánchez-Vázquez, F. J. (2010). Molecular cloning, tissue distribution, and daily rhythms of expression of per1 gene in European sea bass (Dicentrarchus labrax). Chronobiol. Int. 27, 19–33.

Sánchez-Vázquez, F. J., Madrid, J. A., Zamora, S. and Tabata, M. (1997). Feeding entrainment of locomotor activity rhythms in the goldfish is mediated by a feeding-entrainable circadian oscillator. J. Comp. Physiol. A 181, 121–132.

Sanogo, Y. O., Band, M., Blatti, C., Sinha, S. and Bell, A. M. (2012). Transcriptional regulation of brain gene expression in response to a territorial intrusion. Proc. R. Soc. B 279, 4929–4938.

Sanogo, Y. O., Hankison, S., Band, M., Obregon, A. and Bell, A. M. (2011). Brain transcriptomic response of threespine sticklebacks to cues of a predator. Brain Behav. Evol. 77, 270–285.

Schmid, B., Helfrich-Förster, C. and Yoshii, T. (2011). A new ImageJ plug-in “ActogramJ” for chronobiological analyses. J. Biol. Rhythms 26, 464–467.

Schwartz, W. J., Helm, B. and Gerkema, M. P. (2017). Wild clocks: preface and glossary. Philos. Trans. R. Soc. Lond. B Biol. Sci. 372, 20170211.

Sevenster, P., Bruijn, E. F.-D. and Huisman, J. J. (1995). Temporal structure in stickleback behaviour. Behaviour 132, 1267–1284.

Sjoberg, K. (1985). Foraging activity patterns in the goosander (Mergus-merganser) and the red-breasted merganser (Mergus-serrator) in relation to patterns of activity in their major prey species. Oecologia 67, 35–39.

Slavík, O. and Horký, P. (2012). Diel dualism in the energy consumption of the European catfish Silurus glanis. J. Fish Biol. 81, 2223–2234.

Sokolove, P. G. and Bushell, W. N. (1978). The chi square periodogram: Its utility for analysis of circadian rhythms. J. Theor. Biol. 72, 131–160.

Takahashi, J. S. (2017). Transcriptional architecture of the mammalian circadian clock. Nat. Rev. Genet. 18, 164–179.

Takahashi, M., Tahara, Y., Tsubosaka, M., Fukazawa, M., Ozaki, M., Iwakami, T., Nakaoka, T. and Shibata, S. (2018). Chronotype and social jetlag influence human circadian clock gene expression. Sci. Rep. 8, 10152.

Tamai, T. K., Young, L. C. and Whitmore, D. (2007). Light signaling to the zebrafish circadian clock by Cryptochrome 1a. Proc. Natl. Acad. Sci. U. S. A. 104, 14712–14717.

Teboul, M., Barrat-Petit, M. A., Li, X. M., Claustrat, B., Formento, J. L., Delaunay, F., Lévi, F. and Milano, G. (2005). Atypical patterns of circadian clock gene expression in human peripheral blood mononuclear cells. J. Mol. Med. 83, 693–9.

Tessier, I., Boisclair, J., Pomerleau, R. and Thibault, A. (2008). État des connaissances-Parc national du Lac-Témiscouata. QC, Canada: Bibliothèque et Archives nationales du Québec.

Tudorache, C., Slabbekoorn, H., Robbers, Y., Hin, E., Meijer, J. H., Spaink, H. P. and Schaaf, M. J. M. (2018). Biological clock function is linked to proactive and reactive personality types. BMC Biol. 16, 148.

Vatine, G., Vallone, D., Appelbaum, L., Mracek, P., Ben-Moshe, Z., Lahiri, K., Gothilf, Y. and Foulkes, N. S. (2009). Light directs zebrafish period2 expression via conserved D and E boxes. PLoS Biol. 7, e1000223.

Vaze, K. M. and Sharma, V. K. (2013). On the adaptive significance of circadian clocks for their owners. Chronobiol. Int. 30, 413–33.

Vera, L. M., Cairns, L., Sánchez-Vázquez, F. J. and Migaud, H. (2009). Circadian rhythms of locomotor activity in the Nile tilapia Oreochromis niloticus. Chronobiol. Int. 26, 666–681.

Vera, L. M., Madrid, J. A. and Sánchez-Vázquez, F. J. (2006). Locomotor, feeding and melatonin daily rhythms in sharpsnout seabream (Diplodus puntazzo). Physiol. Behav. 88, 167–172.

Vera, L. M., Negrini, P., Zagatti, C., Frigato, E., Sánchez-Vázquez, F. J. and Bertolucci, C. (2013). Light and feeding entrainment of the molecular circadian clock in a marine teleost (Sparus aurata). Chronobiol. Int. 30, 649–661.

Wang, H. (2008a). Comparative analysis of period genes in teleost fish genomes. J. Mol. Evol. 67, 29–40.

Wang, H. (2008b). Comparative analysis of teleost fish genomes reveals preservation of different ancient clock duplicates in different fishes. Mar. Genom. 1, 69–78.

Wang, H. (2009). Comparative genomic analysis of teleost fish bmal genes. Genetica 136, 149–161.

Wang, M., Zhong, Z., Zhong, Y., Zhang, W. and Wang, H. (2015). The zebrafish period2 protein positively regulates the circadian clock through mediation of retinoic acid receptor (RAR)-related orphan receptor α (Rorα). J. Biol. Chem. 290, 4367–4382.

Ware, J. V., Nelson, O. L., Robbins, C. T. and Jansen, H. T. (2012). Temporal organization of activity in the brown bear (Ursus arctos): roles of circadian rhythms, light, and food entrainment. Am. J. Physiol. Regul. Integr. Comp. Physiol. 303, R890–R902.

Wark, A. R., Greenwood, A. K., Taylor, E. M., Yoshida, K. and Peichel, C. L. (2011). Heritable differences in schooling behavior among threespine stickleback populations revealed by a novel assay. PLoS One 6, e18316.

Waterhouse, J., Reilly, T., Atkinson, G. and Edwards, B. (2007). Jet lag: trends and coping strategies. The Lancet 369, 1117–1129.

Weger, M., Weger, B. D., Diotel, N., Rastegar, S., Hirota, T., Kay, S. A., Strähle, U. and Dickmeis, T. (2013). Real-time in vivo monitoring of circadian E-box enhancer activity: A robust and sensitive zebrafish reporter line for developmental, chemical and neural biology of the circadian clock. Dev. Biol. 380, 259–273.

Whitmore, D., Foulkes, N. S. and Sassone-Corsi, P. (2000). Light acts directly on organs and cells in culture to set the vertebrate circadian clock. Nature 404, 87–91.

Whitmore, D., Foulkes, N. S., Strähle, U. and Sassone-Corsi, P. (1998). Zebrafish Clock rhythmic expression reveals independent peripheral circadian oscillators. Nat. Neurosci. 1, 701–707.

Worgan, J. P. and Fitzgerald, G. J. (1981). Diel activity and diet of three sympatric sticklebacks in tidal salt marsh pools. Can. J. Zool. 59, 2375–2379.

Závorka, L., Aldvén, D., Näslund, J., Höjesjö, J. and Johnsson, J. I. (2016). Inactive trout come out at night: behavioral variation, circadian activity, and fitness in the wild. Ecology 97, 2223–2231.

